# Hierarchical Bayesian Modeling of the Relationship between Task Related Hemodynamic Responses and Cortical Excitability

**DOI:** 10.1101/2021.10.22.465452

**Authors:** Zhengchen Cai, Giovanni Pellegrino, Jean-Marc Lina, Habib Benali, Christophe Grova

**Author notes:** **Correspondence:** Zhengchen Cai, Multimodal Functional Imaging Lab, Department of Physics and PERFORM Centre, Concordia University. Loyola Campus, Office SP 365.23, 7141 Sherbrooke Street West, Montreal, QC, H4B 1R6, Canada, Corresponding author.

## Abstract

**Background:** Investigating the relationship between task-related hemodynamic responses and cortical excitability is challenging because it requires simultaneous measurement of hemodynamic responses while applying non-invasive brain stimulation. Moreover, cortical excitability and task-related hemodynamic responses are both associated with inter-/intra-subject variability. To reliably assess such a relationship, we applied hierarchical Bayesian modeling.

**Methods:** This study involved 16 healthy subjects who underwent simultaneous Paired Associative Stimulation (PAS10, PAS25, Sham) while monitoring brain activity using functional Near-Infrared Spectroscopy (fNIRS), targeting the primary motor cortex (M1). Cortical excitability was measured by Motor Evoked Potentials (MEPs), and the motor task-related hemodynamic responses were measured using fNIRS 3D reconstructions. We constructed three models to investigate: 1) PAS effects on the M1 excitability; 2) PAS effects on fNIRS hemodynamic responses to a finger tapping task, and 3) the correlation between PAS effects on M1 excitability and PAS effects on task-related hemodynamic responses.

**Results:** Significant increase in cortical excitability was found following PAS25, whereas a small reduction of the cortical excitability was shown after PAS10 and a subtle increase occurred after sham. Both HbO and HbR absolute amplitudes increased after PAS25 and decreased after PAS10. The probability of the positive correlation between modulation of cortical excitability and hemodynamic activity was 0.77 for HbO and 0.79 for HbR.

**Conclusion:** We demonstrated that PAS stimulation modulates task-related cortical hemodynamic responses in addition to M1 excitability. Moreover, the positive correlation between PAS modulations of excitability and hemodynamics brought insight into understanding the fundamental properties of cortical function and cortical excitability.

## 1 Introduction

The association between hemodynamic responses to a task and excitability of the corresponding cortical region helps to understand the relationship between cortical metabolic demand and cortical readiness. This knowledge might be useful to expand the field of application of non-invasive brain stimulation for treating brain disorders in which modulation of hemodynamic activity is desired. Currently, there are several non-invasive brain stimulation techniques capable of interacting with cortical function, that are potentially useful to treat neurological conditions. One of the well-known techniques is repetitive Transcranial Magnetic Stimulation (rTMS), consisting in TMS pulses repeated in a train following a certain frequency to induce Long-Term Potentiation (LTP)-like and Long-Term Depression (LTD)-like plasticity, which is typically measured as changes of cortical excitability [Ridding and Rothwell, 2007; Di Pino et al., 2014]. Another way to induce plasticity relies on the concept of Spike Timing Dependent Plasticity (STDP, Levy and Steward, 1983; Rossini et al., 2015) and is known as Paired Associative Stimulation (PAS) (Mariorenzi et al., 1991; Stefan, 2000). In PAS paradigm, Transcranial Magnetic Stimulation (TMS) over the motor region (M1) excites the pyramidal cells mimicking a postsynaptic spike, whereas somatosensory activations induced by Median Nerve Stimulation (MNS) propagate from the wrist to pyramidal cells of the motor cortex, acting as a presynaptic spike. TMS and MNS are delivered with proper timing to modulate cortical excitability. Around 25ms (PAS25) Interstimulus Interval (ISI) between MNS and TMS increases excitability, whereas 10ms (PAS10) ISI inhibits the primary motor cortex. To measure the level of cortical excitability elicited by such a technique, the peak-to-peak amplitude of Motor Evoked Potentials (MEPs) measured by electromyography (EMG) on the hand muscle while delivering single pulse TMS (spTMS) to the primary motor cortex, is usually considered [Suppa et al., 2017].

The effects of non-invasive brain stimulation are not limited to cortical excitability, but also involve brain metabolism. In animal studies using invasive optical imaging, Allen et al., (2007) demonstrated that low-frequency repetitive rTMS applied to the cat’s visual cortex, a paradigm known for reducing cortical excitability, induced an immediate increase of tissue oxygenation followed by a prolonged reduction of oxygenation lasting approximately two minutes. More recently, Seewoo et al., 2019 conducted a study on rats combining functional Magnetic Resonance Imaging (fMRI) and proton Magnetic Resonance Spectroscopy (MRS) and showed: 1) increases in resting-state connectivity (e.g., Interoceptive/default mode network, cortico-striatal-thalamic network and Basal ganglia network), GABA, glutamine and glutamate levels following high-frequency rTMS (known for increasing cortical excitability) and 2) Reduced connectivity and glutamine levels after low-frequency rTMS stimulations.

In human studies, similar investigations are more challenging since they require the combination of non-invasive neuroimaging and non-invasive brain stimulation approaches. For instance, fMRI [Kwong et al., 1992; Bandettini et al., 1992; Glover, 2011] is a widely used modality to measure the hemodynamic activity, and could be exploited to assess the hemodynamic fluctuations related to TMS interventions [Navarro De Lara et al., 2015; Tik et al., 2017]. However, simultaneous fMRI/TMS acquisition is challenging, since it requires specific MRI and TMS coils and fMRI sequences [Navarro De Lara et al., 2015; Wang et al., 2017]. This is the main reason why most studies conducted fMRI sessions before and after TMS interventions, following a so-called offline approach [Siebner et al., 2009].

Alternatively, functional Near-InfraRed Spectroscopy (fNIRS) non-invasively measures fluctuations of both oxygenated- and deoxygenated-hemoglobin concentration changes in the human cortex (i.e., HbO and HbR), usually offering better temporal resolution than fMRI [Jöbsis, 1977; Scholkmann et al., 2014]. fNIRS relies on the optical signal, which is not sensitive to fluctuations of electromagnetic fields, as opposed to the fMRI signal. Therefore, fNIRS offers better compatibility for simultaneous acquisition during TMS [Curtin et al., 2019]. In Cai et al., 2022b, we demonstrated that simultaneous PAS-fNIRS is feasible and investigated changes in excitability and hemodynamic activity due to non-invasive neuromodulation. Among the many non-invasive brain stimulation techniques, we focused our attention on PAS, due to methodological and biological reasons. First, a previous simultaneous TMS/fNIRS study reported that physiological fluctuations of respiration and heart rate are largely influenced by trains of TMS pulses [Näsi et al., 2011]. In other words, TMS trains may influence both systemic hemodynamic (scalp signal) and cortical hemodynamic responses. Second, the frequency of stimulation pairs in PAS is 0.1Hz or less [Suppa et al., 2017], therefore PAS intervention is likely to introduce significantly less or even no systemic physiological fluctuations when compared to rTMS. Third, PAS allows multiple effects (excitation, inhibition, sham-no effect) by simply changing the interstimulus interval between MNS and TMS, thus offering similar duration and frequency of stimulations across conditions, allowing a more balanced experimental design.

All neuromodulatory techniques are known to produce variable results [Ridding and Ziemann, 2010]. This aspect goes beyond the technical difficulties and challenges encountered when studying the relationship between excitability and hemodynamic activity and involves both intra- and inter-subject variability inherent to the effects of brain stimulation and hemodynamic activity associated with a task. The ability of PAS in eliciting significant changes in cortical excitability has been replicated by several studies [Stefan, 2000; Wolters et al., 2005; Tsang et al., 2015; Lee et al., 2017; Suppa et al., 2017], but some studies have observed that only 39% of subjects showed expected MEP amplitude increase after conducting PAS25 [López-Alonso et al., 2014]. Similar PAS efficiency (lower than 50%) has been suggested in a review study [Suppa et al., 2017]. Inter-subject variability of the task-evoked hemodynamic response has also been reported, whether measured using fNIRI [Witt et al., 2008] or fNIRS [Novi et al., 2020].

These variability issues may explain negative findings when reporting the correlation between cortical excitability and hemodynamic activities. For instance, Kriváneková et al., (2013) investigated the relationship between the primary motor cortex (M1) excitability and Blood-Oxygen-Level-Dependent (BOLD) signal using PAS stimulation and “offline” fMRI acquisitions. They reported no significant correlation between PAS effects on task-related BOLD response and its effects on M1 excitability. In our previous study [Cai et al., 2022b], using fNIRS and PAS we first found a significant positive correlation between fluctuations of cortical excitability (represented by MEP amplitude) and the fluctuations of HbO activity. However, when further investigating the relationship between PAS effects on task-related HbO/HbR changes (estimated using the HbO or HbR ratio, calculated as the post-over pre-intervention amplitudes) and its effects on M1 excitability (estimated using the ratio of MEP, calculated as the post-over pre-intervention amplitudes), we also found no significant correlations. Therefore, it seems essential to carefully take into account the intrinsic variability of both cortical excitabilities elicited by non-invasive brain stimulation and hemodynamic responses to tasks, when investigating such correlations. Note that this is not a limitation specific to PAS, but rather an inherent issue of all neuromodulatory techniques and depends on several factors including individual anatomy, genetic susceptibility, hormonal activity, the position of the coil for stimulation, and circadian fluctuations of excitability. While much of the variability is inherent and non-avoidable, it is however possible to properly model them. This could be achieved by applying Bayesian modeling.

To conduct accurate and robust investigations, we propose to incorporate the variability of data using a hierarchical Bayesian model to assess PAS effects on both cortical excitability and hemodynamic responses, as well as the correlation between these two measures.

Hierarchical Bayesian modeling allows taking into account heterogeneity of the variables of interest (i.e., MEP, HbO/HbR) at each level (i.e. intra-subject, inter-subject, intervention type: PAS25, PAS10, sham) of the analysis [Papaspiliopoulos et al., 2007; Betancourt and Girolami, 2015]. Moreover, when considering a hierarchical structure, partial pooling can reduce the uncertainty of estimated parameters [Gelman et al., 2013; McElreath, 2020]. This means that group-level and individual-level estimations could inform each other to regularize the estimation of the uncertainty of each parameter. Moreover, Bayesian inference allows estimating the statistical expectation of each parameter of the model by sampling the joint posterior distributions. Thanks to the developments in Bayesian data analysis workflow during the last decade, Bayesian inference has become more accessible and can provide accurate and reliable estimations of the posterior distribution. For instance, the most recent implementations of the Hamiltonian Monte Carlo (HMC) algorithm [Duane et al., 1987] called the dynamic HMC [Betancourt, 2017; Betancourt, 2019] is available as an open-source Bayesian statistical modeling and computation platform called Stan [Stan Development Team, 2020a]. This technique not only accurately and efficiently samples the joint posterior distribution, but also provides robust estimations by quantitatively diagnosing pathological behaviors of Markov Chain Monte Carlo (MCMC) chains that are used to sample the joint posterior distributions [Betancourt and Girolami, 2015; Betancourt, 2017].

Considering the above advantages of the Bayesian approach, in this study, we applied a Bayesian data analysis workflow [Gabry et al., 2019; Gelman et al., 2020b] on our TMS/fNIRS dataset to further investigate the relationship between the PAS effects on task-related hemodynamic responses and its effects on M1 excitability, for which we reported small effects using conventional analysis in Cai et al., 2022b. Bayesian inference allows us to handle the intrinsic variability of the data. Moreover, it will also offer the unique opportunity to apply posterior predictive simulations [Gabry et al., 2019; Gelman et al., 2020b] to investigate new questions that could not be addressed using conventional analysis. We hypothesize that enhanced brain excitability should be associated with higher hemodynamic activity elicited by a finger tapping task, and decreased excitability should be associated with a reduced hemodynamic response to the task. In this work, we first summarize the study design, data acquisition, and data preprocessing prior to the Bayesian framework. Then, we proposed three hierarchical Bayesian models: Model#1: PAS effects on M1 excitability measured using MEP; Model#2: PAS effects on task-related hemodynamic responses measured using fNIRS, and Model#3: correlation between PAS modulated excitability changes and PAS modulated hemodynamic changes. The variability of each measurement was carefully considered in each model and at each level to conduct reliable estimations of the intervention effects and correlations. Statistical inferences were made via posterior predictive simulations [McElreath, 2020]. Diagnostics of the models were conducted to ensure the robustness of the estimated posterior distributions.

## 2 Material and methods

### 2.1 Study design and subjects

Nineteen subjects (19 – 35 years old, male and right-handed) with no history of neurological disorders and no medications acting on the central nervous system were selected to participate in the study. We only included male participants in order to minimize the confounding of cortical excitability changes due to the menstrual cycle [Hattemer et al., 2007; Lee et al., 2017]. This study was approved by the Central Committee of Research Ethics of the Minister of Health and Social Services Research Ethics Board (CCER), Québec, Canada. All subjects signed written informed consent prior to participation. They also underwent a screening procedure to confirm no contraindications to MRI or TMS [Rossi et al., 2009; Suppa et al., 2017]. Subjects were instructed to have a regular sleep cycle for the days and not to take caffeine for at least 90 minutes before the data acquisition.

The experiment paradigm of this study is illustrated in Fig.1.a. Three different intervention sessions were performed at least two days apart to minimize carryover effects. Experimental sessions were performed in a pseudorandomized order, each session consisted of five time-ordered sections, defined as follows:

1. ***A block designed finger-tapping task composed of 20 blocks, 10s of finger-tapping followed by 30s ~ 60s of resting was conducted within each block.*** Subjects were asked to tap their left thumb to the other 4 digits sequentially around 2Hz (Fig.1.a1). This long-range jitter was designed to prevent the task responses from phase locking to the undergoing physiological hemodynamic oscillations [Aarabi et al., 2017], therefore, reducing the physiological confounding on the task-related response at the stage of experiment paradigm design. Tapping onsets/offsets were instructed by auditory cues.
2. ***An event-related designed single pulse TMS (spTMS) composed of 75 events, jittered from 5s to 25s (Fig.1.a2).*** TMS procedures were performed with neuronavigation (Brainsight neuro-navigation system - Rogue-Research Inc, Canada) and based on subject-specific anatomical MRI. TMS was delivered with a figure-8 coil (Magstim double 70mm remote control coil) connected to a Magstim 200^2^ stimulator (Magstim Company, U.K.). In order to target M1, the coil was placed tangentially to the scalp and with a 45° angle to the midline of the head (Fig.1.c), so to maximize stimulation efficiency [Thomson et al., 2013; Raffin et al., 2015a]. The individual ‘hot spot’ was defined for each session as the location with the largest Motor Evoked Potentials (MEPs) amplitude measured on the left thumb (Abductor Pollicis Brevis, APB) using electromyography (EMG). Stimulation intensity was set to 120% of the resting motor threshold (RMT), defined according to the maximum-likelihood parameter estimation by a sequential testing approach using MTAT 2.0 (http://www.clinicalresearcher.org/software.html) [Awiszus et al., 1999; Ah Sen et al., 2017]. All TMS procedures followed the recommendations of the International Federation of Clinical Neurophysiology [Rossi et al., 2009] and no participants reported any considerable discomfort or side effects.
3. ***A PAS session to modulate the M1 cortical excitability (Fig.1.a3).*** PAS intervention consisted either in PAS25 to increase excitability, PAS10 to decrease excitability or sham-PAS. PAS was conducted with 100 pairs of electrical median nerve stimulation (MNS) on the left wrist, followed by TMS pulse delivered over the right M1, with a fixed interval of 10s between paired stimulations, for a total intervention of 18 minutes, as suggested in Suppa et al., (2017). MNS was delivered with a Digitimer (Digitimer DS7A, U.K) at the left median nerve and with an intensity equal to 300% of the subject-specific perceptual threshold, as suggested in Stefan, 2000. TMS intensity was the same as for spTMS paradigm, i.e., 120% of RMT. After estimating subject-specific N20 response to electrical MNS using bipolar electroencephalogram (EEG) (BrainAmp ExG, Brain Products GmbH, Germany) on CP3 and CP4 electrodes, the interstimulus intervals (ISI) between MNS and TMS were determined based on individual N20 values: as N20+5ms for PAS25 and N20-5ms for PAS10 [Carson and Kennedy, 2013]. Sham parameters (e.g., MNS intensity, coil position, ISI) were the same as PAS25, but TMS was not delivered, and instead, its sound (‘TMS click’) was played via a stereo speaker.
4. ***Repetition of the event-related designed spTMS (Fig.1.a4) after the intervention.*** By comparing the MEPs measured during pre-intervention and post-intervention sessions, PAS intervention effects on M1 cortical excitability could be assessed.
5. ***Repetition of the block designed finger tapping task (Fig.1.a5) after the intervention.*** Similarly, the corresponding effects on task-evoked hemodynamic responses could be assessed by comparing HbO/HbR concentration changes measured during preintervention and post-intervention sessions.

**Fig.1.**
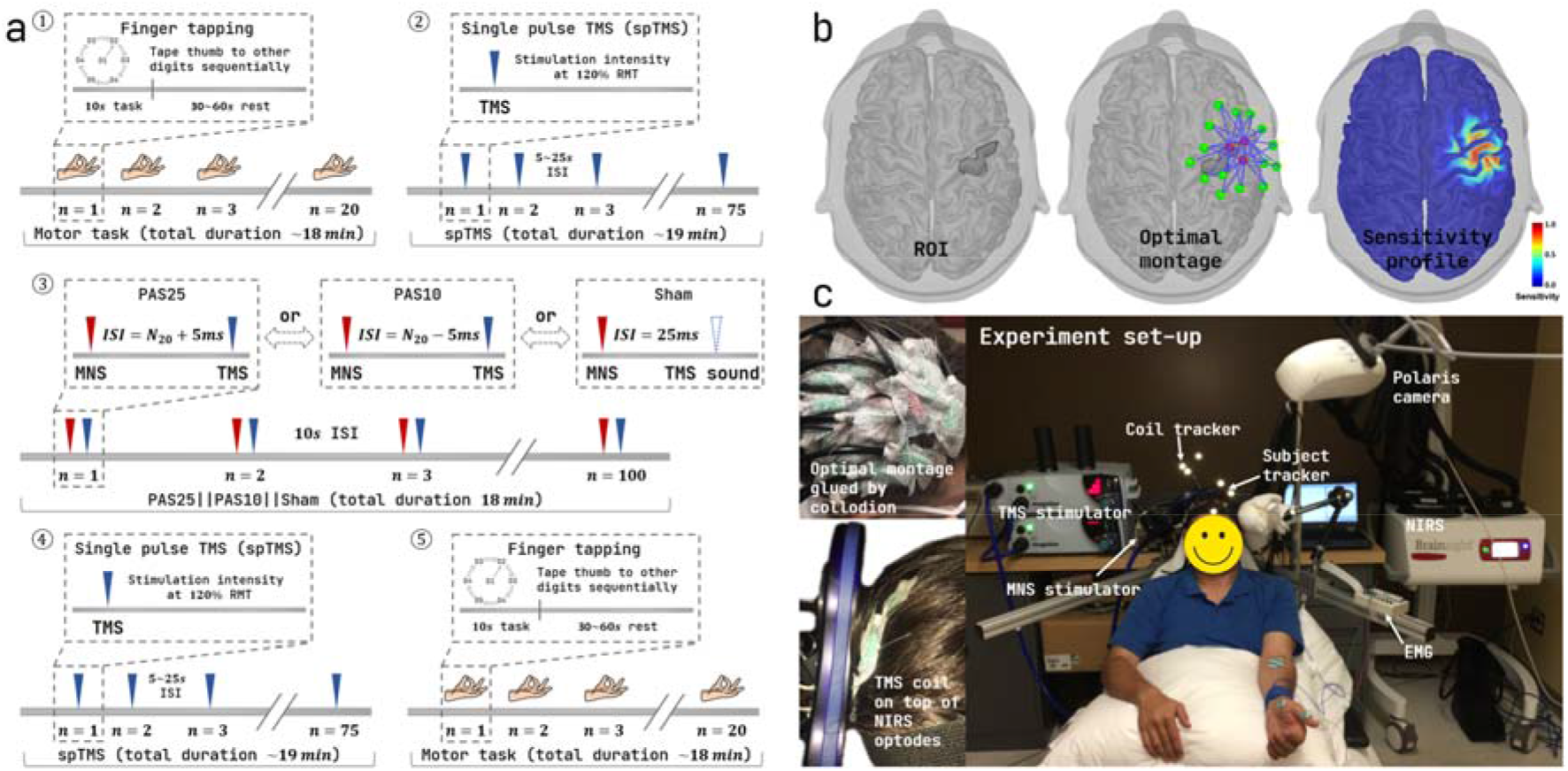
Experimental paradigm and set-up. a) experimental paradigm ordered by time: 1) a block designed finger-tapping task consisted of 20 blocks, each contained 10s task and 30s to 60s rest; subjects were informed to tap their left thumb to the other 4 digits sequentially at around 2Hz; 2) an event-related designed single pulse TMS (spTMS) run consisted of 75 events jittered from 5s to 25s. 3) PAS25/PAS10/sham-PAS consisted of 100 pairs of stimulations, interleaved by 10s; 4) and 5) repeated 2) and 1), respectively, after the PAS intervention. b) personalized optimal montage for fNIRS acquisition. 3 sources (red dots) and 15 detectors (green dots) were selected to optimize the sensitivity of fNIRS montage to a predefined ROI, the right M1 hand knob (outlined using a black profile) along the cortical surface. c) an overview of the experimental set-up, the personalized optimal montage was glued on the scalp using clinical adhesive – collodion; TMS coil was placed on top of the fNIRS optodes to target the ‘hot spot’ which corresponded to subject’s left thumb, note that the low-profile feature of the fNIRS optodes allowed less TMS intensity decreases when departing from the scalp surface; a neuro-navigation system was used to guide the placement of the TMS coil and the digitization of the fNIRS optodes.

### 2.2 Data acquisitions

#### 2.2.1 Anatomical MRI

Individual anatomical MRI was acquired to guide TMS and to calculate the head model required for fNIRS acquisition planning and fNIRS reconstructions. A General Electric Discovery MR750 3T scanner at the PERFORM Center of Concordia University, Montréal, Canada, was used to scan: 1) T1-weighted images using the 3D BRAVO sequence (1 × 1 × 1 mm^3^, 192 axial slices, 256 × 256 matrix) and 2) T2-weighted images using the 3D Cube T2 sequence (1 × 1 × 1 mm voxels, 168 sagittal slices, 256 × 256 matrix).

#### 2.2.2 Motor Evoked Potentials

MEPs induced by spTMS pulses were measured to assess the M1 cortical excitability. A BrainAmp ExG bipolar system (BrainAmp ExG, Brain Products GmbH, Germany) was used to record EMG of the right abductor pollicis brevis (APB) muscle, with 2 TECA disposable 20mm disk electromyography (EMG) electrodes attached with a standard belly-tendon montage (Fig1.c).

#### 2.2.3 Functional Near-Infrared Spectroscopy

fNIRS data were acquired to estimate the finger-tapping evoked hemodynamic responses (i.e., HbO/HbR responses). fNIRS data were acquired at 10Hz using a Brainsight fNIRS system (Rogue-Research Inc, Canada) with two wavelengths – 685nm and 830nm. fNIRS optodes were placed on the subject’s scalp using a clinical adhesive called collodion (Fig.1.c) to reduce motion artifacts [Yücel et al., 2014; Pellegrino et al., 2016; Machado et al., 2018] and to ensure a better contact with the skin when compared to standard fNIRS caps. A personalized optimal montage developed by our group [Machado et al., 2014; Pellegrino et al., 2016; Machado et al., 2018; Cai et al., 2021] was used to maximize the sensitivity of fNIRS channels to a predefined region of interest (ROI) - the individual ‘hand knob’ region (see Fig.1.b) manually defined along the right M1 cortical surface which controls the left hand movement [Raffin et al., 2015b]. The resulted personalized optimal montage consisted of 3 sources and 15 detectors (see Fig.1.b). The distance between each source-detector pair was constrained to range from 2.0 cm to 4.5 cm. Each source was positioned to construct at least 13 channels among the 15 detectors ensuring a high spatial overlap between channels, to allow accurate local reconstruction along the cortical surface [Cai et al., 2021; Cai et al., 2022a]. A proximity detector was added at the center of 3 sources to record the physiological hemodynamics fluctuations within the scalp. A Brainsight neuro-navigation system coregistered with the subject’s specific T1 MRI was used to guide the installation and to digitize the position of fNIRS sources and detectors glued at their optimal positions. Additional 150 points were digitized on the head surface to allow accurate montage registration with the anatomical MRI, as a prerequisite for computing the fNIRS forward model. fNIRS data were acquired continuously during the whole experimental session, as described in Fig.1.a.

From the nineteen subjects selected for this study, one was excluded due to low sensitivity to TMS, and two were excluded because they exhibited poor fNIRS signal qualities. Four subjects dropped out after the first session due to personal reasons, resulting in 16 PAS25, 12 PAS10 and 12 sham sessions. Please note that starting from here, we will denote: 1) as ***“Session”,*** one specific acquisition, consisting in any PAS intervention type, of one subject including experiments 1 to 5 illustrated in Fig.1.a (e.g., PAS25 for Sub01); as ***“Run”,*** one specific experiment (spTMS or finger tapping), before or after any PAS intervention type, for one subject (e.g., pre-PAS25 spTMS for Sub01); and as ***“Time”,*** the differentiation whether one specific experiment was conducted before or after PAS intervention (e.g., pre-PAS25 vs. post-PAS25).

### 2.3 Data preprocessing

#### 2.3.1 EMG data processing

EMG data collected during spTMS runs were processed using Brainstorm software [Tadel et al., 2011] (https://neuroimage.usc.edu/brainstorm/) to extract MEP amplitudes. Raw EMG data were first band-pass filtered between 3 and 2000Hz. A time window from −10ms to 100ms around the stimulation onset was defined to extract MEP trials. These trials were then baseline corrected (−10ms to 0ms), and the peak-to-peak amplitude of each MEP trial was calculated. Note that throughout the analysis reported in this study, none of the single MEP trials was excluded to preserve the intrinsic variability of MEP peak-to-peak amplitude measures. Hereby, for convenience, we will denote as “MEP”, the actual MEP peak-to-peak amplitude, as usually considered in TMS literature [Stefan, 2000; Wolters et al., 2005; Tsang et al., 2015; Lee et al., 2017; Suppa et al., 2017].

The output of the whole EMG data preprocessing section was a set of 75 MEPs estimated for each participant (specified by subject ID from 1 to 16), each intervention (PAS25, PAS10 or sham) and time (pre-PAS or post-PAS).

#### 2.3.2 fNIRS data processing

fNIRS data processing was performed using the open-source fNIRS processing plugin - NIRSTORM (https://github.com/Nirstorm/nirstorm) developed in our lab within Brainstorm software environment [Tadel et al., 2011] (https://neuroimage.usc.edu/brainstorm/). Raw fNIRS data were first preprocessed following standard recommendations [Yücel et al., 2021] and then converted to optical density changes (i.e., ΔOD). For each task run, 20 ΔOD epochs were extracted within a time window ranging from −10s to 30s around task onsets. To reduce motion artifacts and obtain the distribution of averaged ΔOD epochs for each run, we sub-averaged 16 out of 20 ΔOD epochs for all possible unique combinations (i.e., 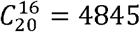 possibilities) and selected 50 of these sub-averaged ΔOD epochs that were ranging below and above the median of the signal to noise ratio (SNR) of all sub-averaged trials, therefore resulting in 101 sub-averaged ΔODs. This approach was to exclude eventual motion artifacts contaminated epochs while preserving the variability of task-evoked fNIRS signal changes in each run [Cai et al., 2022b]. To obtain the distribution of spatiotemporal map of HbO/HbR responses for each finger-tapping run along the cortical surface, we applied a 3D fNIRS reconstruction workflow [Cai et al., 2021; Cai et al., 2022a] using personalized optimal montage and maximum entropy on the mean (MEM) to these 101 sub-averaged ΔODs. The resulting HbO/HbR spatiotemporal maps of each subject during each finger-tapping run (e.g., 101 HbO maps for Sub01 during pre-PAS25 finger-tapping) were finally co-registered to the mid-surface of the MNI ICBM152 template [Fonov et al., 2009; Fonov et al., 2011], using FreeSurfer spherical transformation. A region of interest (ROI) was defined along the template surface as the “hand knob”, to cover the cortical regions that control finger tapping. Finally, reconstructed HbO/HbR time courses (0s to 30s) within this “hand knob” ROI were averaged to represent the hemodynamic responses of each specific finger-tapping run. The output of the whole fNIRS data preprocessing section was a set of 80 runs: 40 Sessions (16 PAS25+12 PAS10+12 sham) × 2 Times (pre- and post-intervention) of 101 reconstructed HbO/HbR time course, for each run specified by subject (ID 1 to 16), intervention (PAS25, PAS10 or sham) and time (pre-PAS or post-PAS). Further details describing fNIRS data processing are provided in Appendix 1.

### 2.4 Hierarchical Bayesian Modeling

For the notation in the following model equations, we used small letters to denote a variable (e.g., *μ* for the mean of a Gaussian distribution) and capital letters to denote a matrix (e.g., Σ for the covariance matrix of a multivariate Gaussian distribution). A list of values from one specific variable is represented by a small letter along with a subscript letter, for instance, a symbol *μ_s_* refers to a list of means, and the subscript *s* represents each individual element of this list (mean for the *s^th^* session). The dimensionality of each list is given by the range of *s* (e.g., *s* = 1,2,3,…4, *for s^th^ session*). If the subscript letter is contained in square brackets, it means that the elements in this list variable are differentiated by the model using index variables. For instance, *i* in *intercept*_[*i*=1,2,3]_ indicates our model differentiates the intercept parameter for each intervention type by index variable *i* = 1,2,3, 1 for PAS25, 2 for PAS10, and 3 for sham.

#### 2.4.1 Hierarchical Bayesian Model #1: Assessment of PAS effects on cortical excitability

We proposed a first hierarchical Bayesian model to assess PAS effects on M1 cortical excitability (Fig.2), which was evaluated using the MEPs measured during spTMS runs before (pre-) and after (post-) each PAS intervention. This model consists of two parts: 1) a measurement error model taking into account the variability of MEPs within each spTMS run and 2) a hierarchical model describing post-intervention MEP as a linear function of preintervention MEP.

**Fig.2.**
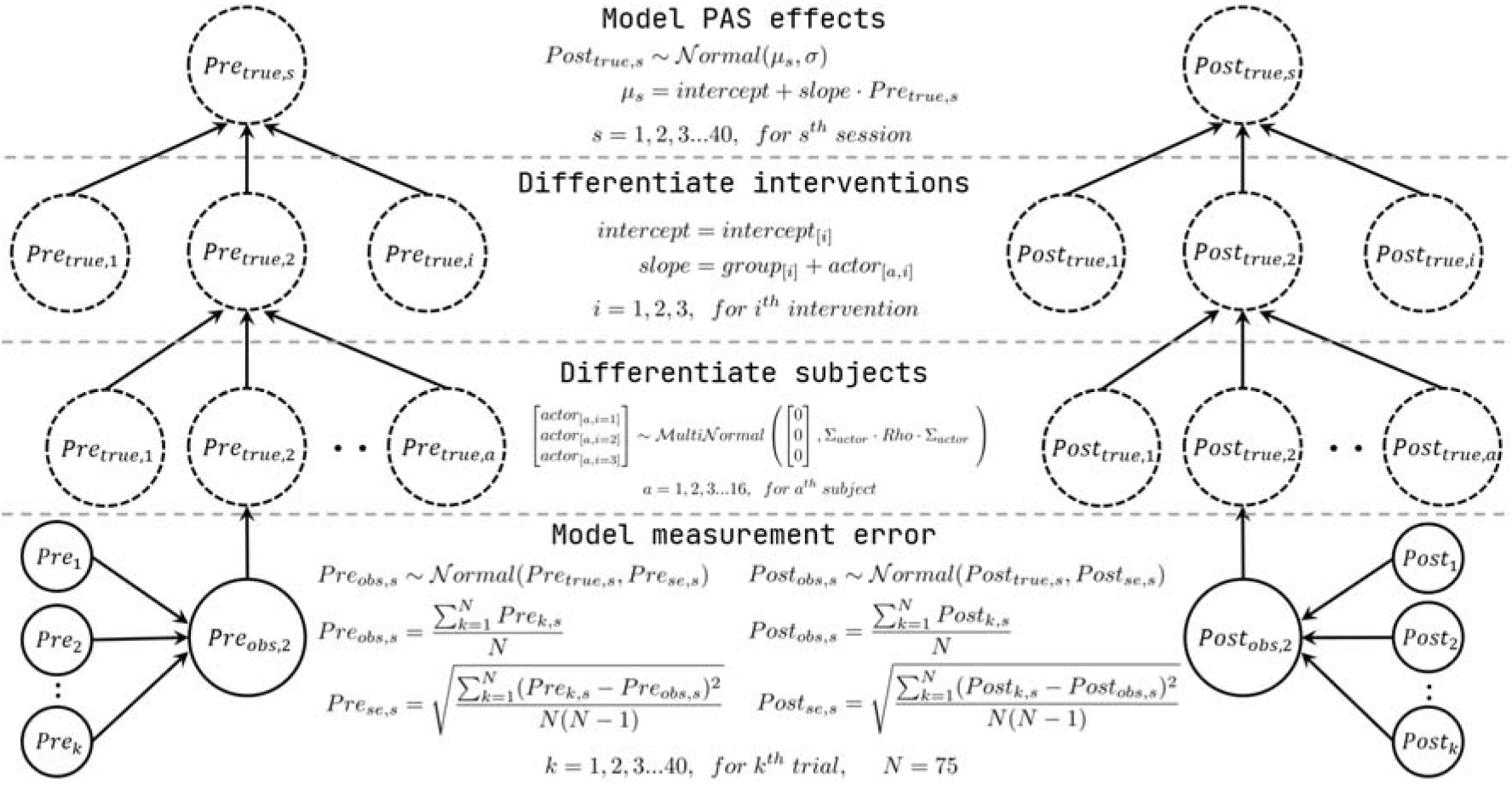
The hierarchical model of PAS effects on either cortical excitability (MEP) or hemodynamic responses (Hb). From bottom to top, 1) a measurement error model assuming the mean of the variable of interest (either the observed MEPs, e.g., 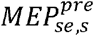 for Model #1, or a spline weight of HbO/HbR time course, e.g., 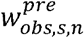 for Model #2) for each run at a different time (pre-/post-) was drawn from a Gaussian distribution. The mean of this Gaussian distribution is the ‘true’ value of the variable of interest, and the scale is the corresponding standard error; 2) each subject and 3) intervention were differentiated using index variables; 4) PAS effects were modeled by linear regression in which the ‘true’ post-variable of interest was predicted by the ‘true’ pre-variable of interest. Solving this hierarchical model by Bayesian allows partial pooling on each parameter to reduce the uncertainty.

##### 1) A model of measurement error

We assume the “empirical” mean of the observed MEP in each run to be drawn from a Gaussian distribution with the mean equal to the ‘true’ MEP amplitude and the scale equal to the standard error of all MEPs trials. The ‘true’ and observed (‘obs’) MEP of the pre-PAS spTMS run can be expressed as follows,

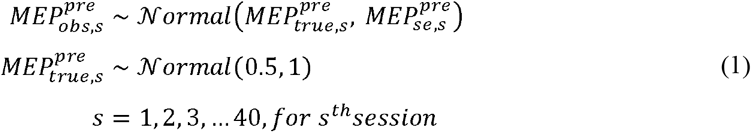

where 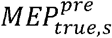 is the ‘true’ value of the pre-PAS mean MEP for session *s*. 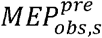 is the “empirical” mean of the observed MEP from the same run expressed as,

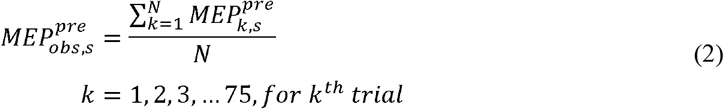

where 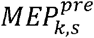 represents the pre-PAS MEP of the *k^th^* trial from a total of N=75 trials in session *s*. The corresponding measurement error 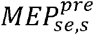 is then represented by the standard error of the MEP over all 75 trials, estimated as in equation (3). Note that we considered here the standard error of MEP samples to represent the scale of the population distribution in equation (1),

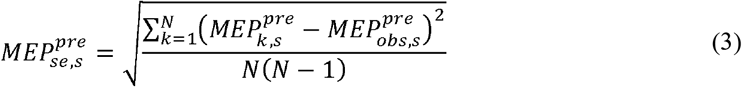

Finally, substituting (2) and (3) into (1), both empirical mean and variance estimated over the 75 observed pre-PAS MEPs of a specific session were modeled to estimate the ‘true’ corresponding amplitude. For the prior distribution of 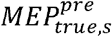, we applied a weakly informed prior [Gelman et al., 2008; Gelman et al., 2017; Gabry et al., 2019] consisting in a Gaussian distribution 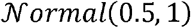. Note that all observed MEPs (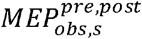 and 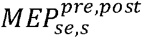, for 40 *Sessions* × 2 *Times* (*pre and post PAS*) = 80) were normalized by the global maximum value of 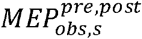 to ensure MEP values were covering the [0,1] range. Therefore, 0.5 appeared as an appropriate prior of the mean when nothing is known about the MEP amplitude, except the range (i.e., (0 + 1)/2 = 0.5).

The ‘true’ MEP in the post-PAS spTMS run was then modeled as follows,

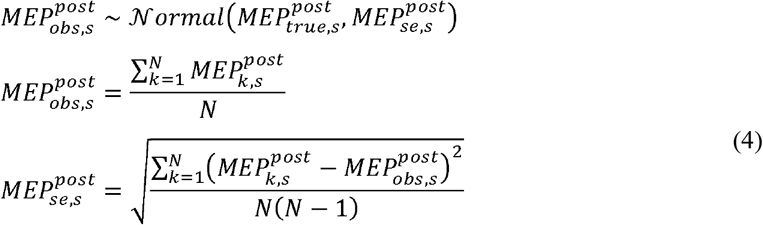

Note that the prior distribution of the parameter 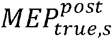 is defined as a function of pre-PAS MEP measures in the next section, within the context of hierarchical linear regression.

##### 2) Hierarchical linear regression

PAS effects on M1 cortical excitability were then modeled using a linear regression model, in which 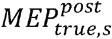 and 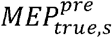 were considered as the dependent and predictor variables, respectively.

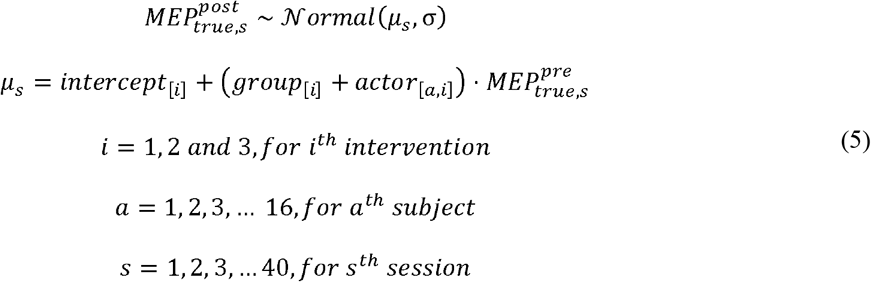

where *μ_s_* is the mean of 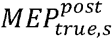, predicted by 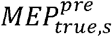 using the following linear model: 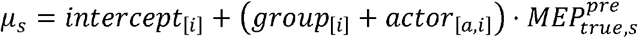, and σ is the error of the linear regression (i.e., the scale of this normal distribution). We added the following index variables to differentiate subject, intervention (PAS25/PAS10/sham) and time (pre-/post-PAS) in the model. *intercept*_[*i*]_ is the intercept of the linear regression, for the *i^th^ intervention,* in which *i* = 1, 2 *and* 3 refers to PAS25, PAS10 and sham, respectively. The slope parameter is modeled using two parts, a group-level slope parameter *groups*, specific for each intervention *i*, and a parameter modeling inter-subject variability, denoted as *actor*_[*a,i*]_, for each intervention *i* and each subject *a*, associated with the following prior model:

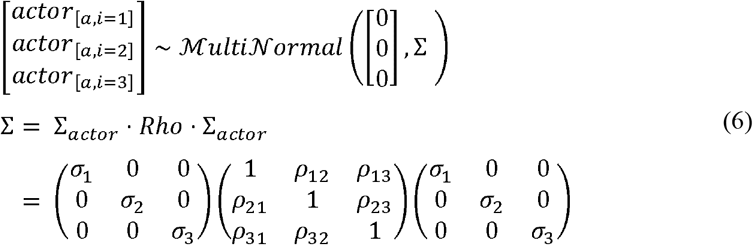

where 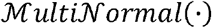 is a multivariate Gaussian distribution to model the interaction, which allows the effects of each specific intervention to vary for each subject, meaning each subject can respond to each intervention differently. We defined this multivariate Gaussian prior distribution to have zero means (3 elements vector) therefore assuming all the subjects to have zero mean deviation around the group-level slope parameter *group*_[*i*]_. *Rho* and *∑_actor_* denote respectively the correlation matrix and the scale matrix of the covariance matrix Σ of this multivariate Gaussian distribution. *σ*_1,2,3_ is the scale among all subjects within each intervention group, for example, *σ_3_* is the scale of the vector *actor*_[*a,i*=3]_ for sham. *ρ* is the correlation between pair-wised interventions, for instance, *ρ_12_* represents the correlation between *actor*_[*a,i*=1]_ for PAS25 and *actor*_[*a,i*=2]_ for PAS10.

Weakly informed priors were assigned to the parameters introduced in equations (5) and (6) as follows,

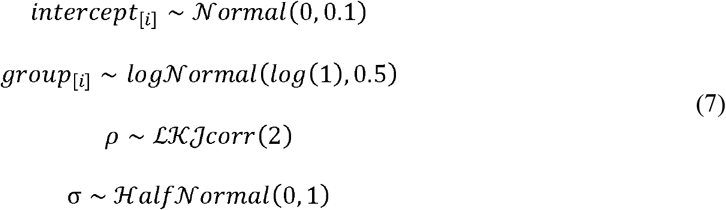

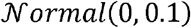 was chosen as prior distribution for *intercept*_[*i*]_ considering that when 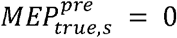, the corresponding 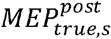 should not be too much apart from 0. 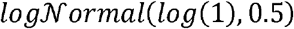 was selected for the group-level slope to ensure a positive value with a median of 1. Therefore, without knowing any intervention type, the slope should be equal to 1, assuming there is no averaged PAS effect among subjects when the intervention type is not known. The Lewandowski-Kurowicka-Joe distribution 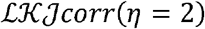, [Lewandowski et al., 2009], was chosen as a weakly informative prior for the correlation parameter *ρ* that does not prioritize extreme correlation values such as ±1, where *η* is a positive parameter. *η* = 1 would denote uniform density of *ρ* from −1 to +1. The larger *η* is (when compared to 1), the least likely the extreme correlation values would occur (sharper probability density distribution). We selected *η* = 2 as a weakly informed prior commonly considered in Bayesian data analysis [McElreath, 2020]. Finally, 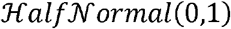 was chosen as the prior distribution for variance parameter to ensure a positive semi-definite value. When variance increase, the corresponding likelihood decreases following the bell shape of the positive half of 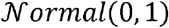 distribution. As denoted previously, we normalized all data 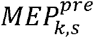 and 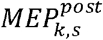 within the range [0,1], therefore 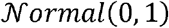 was considered as a conservative (“flat”) prior, when modeling the variance.

#### 2.4.2 Hierarchical Bayesian Model #2: Assessment of PAS effects on task-related hemodynamic responses

A similar hierarchical model was proposed to assess PAS effects on task-related fNIRS hemodynamic responses. This model is very similar to previous Model #1, the main difference being that the input variables to the model are now ‘features’ representing hemodynamic responses to the finger-tapping task. Conventional approaches would extract amplitudes at a specific time sample of HbO/HbR responses (e.g., the hemodynamic peak amplitude), or perform a local average of HbO/HbR responses within a specific time window. However, in this Model #2, we conducted a procedure to model PAS effects over the whole time course of HbO/HbR responses to finger tapping.

To do so, after 3D reconstruction using MEM of all 101 sub-averaged of the fNIRS responses, HbO/HbR time courses were first averaged within the selected M1 ROI in the selected time range [0s, 30s]. To lower the dimension of the input to the model, resulting time courses were projected on B-splines temporal basis functions [Boor, 2001; Gelman et al., 2013; Hastie, 2017]. Therefore, hemodynamic responses were expressed as a weighted linear combination of B-splines basis functions and included as input in our proposed hierarchical Bayesian model (Fig.3). Please note that Hb refers here to either HbO or HbR in the model. The model was fitted separately for each chromophore,

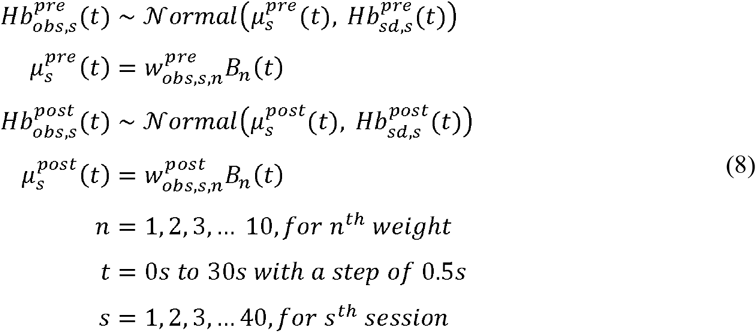

where 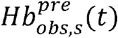 is the observed empirical mean of pre-PAS HbO/HbR responses over all 101 sub-averaged time courses, for a specific finger-tapping run (e.g., finger-tapping run in pre-PAS25 of Sub01) for a specific session *s* at a specific time point *t*. 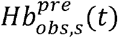 is assumed to follow a Gaussian distribution with a mean of 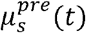 and a scale of 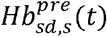, where 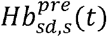 is the corresponding standard deviation estimated over all 101 sub-averaged time courses. Note that all pre- and post-PAS HbO/HbR time courses in one session were normalized by session-specific global maximum absolute amplitude (considering both HbO and HbR) to be within the range [-1,1], since HbR usually exhibits negative amplitude. Then, 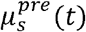 representing the mean time course of the true pre-PAS HbO/HbR for time sample *t* and session *s*, was defined as a linear combination of *n* = 10 B-spline basis functions B_*n*_(*t*) (*n* × *t* matrix) with the corresponding 10 weights 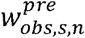 (n-vector). Each basis function *B_n_*(*t*) was defined as a 3^rd^ order polynomial function. A similar model structure was applied to data and parameters when modeling HbO/HbR responses from the post-PAS finger-tapping runs.

**Fig.3.**
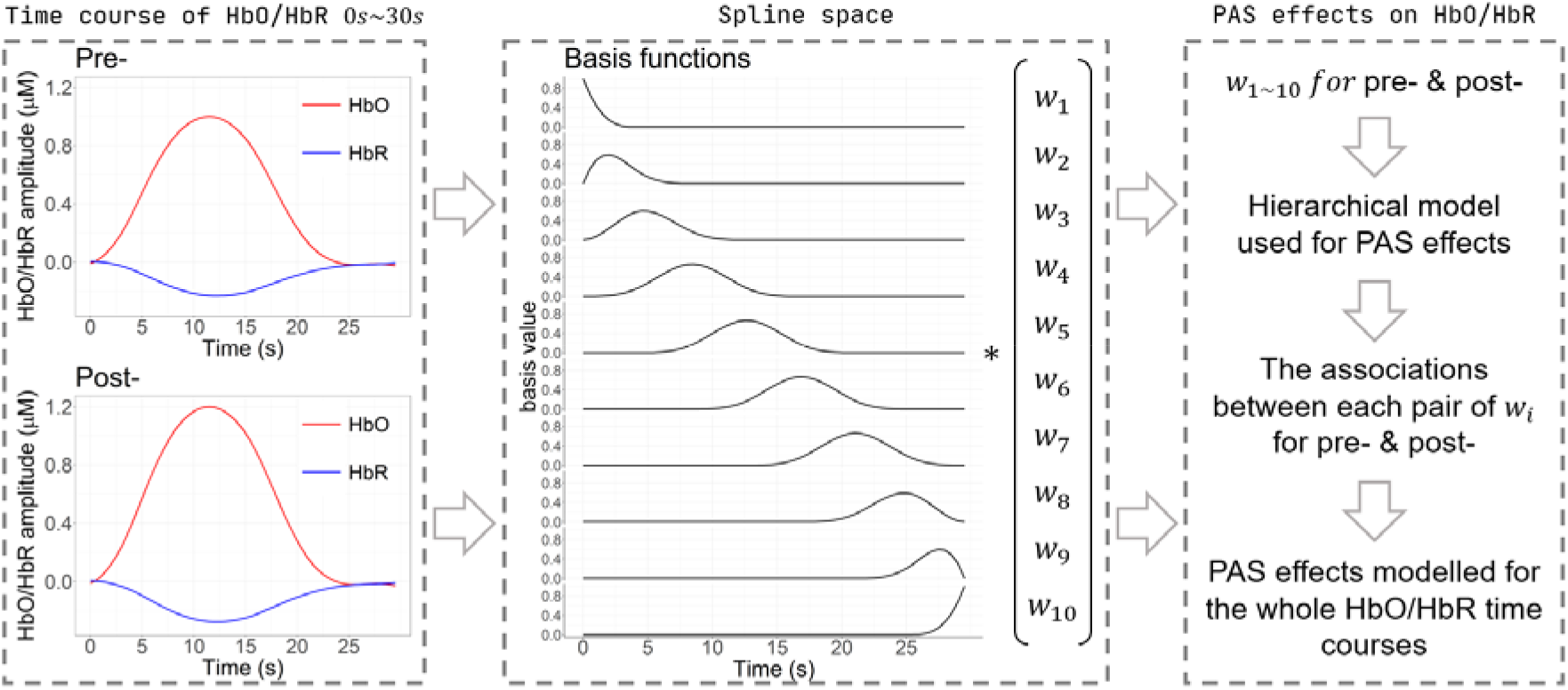
The hierarchical model for PAS effects on the whole HbO/HbR time course. HbO/HbR time courses, from 0s to 30s, before and after each intervention session were selected as the inputs of the model. They were first projected into the spline space which was composed of 10 predefined basis functions (3^rd^ order polynomial with 10 knots). The linear combination of basis function using 10 corresponding weights w_1~10_ was able to fully recover the HbO/HbR time course. The resulting pre- and post-PAS spline weights were then fed into the hierarchical model, similar to Model #1 to estimate the PAS effects on each weight. Therefore, the associations between each pair of weights encapsulated the PAS effects over the whole time course of HbO/HbR hemodynamic responses.

To model the temporal hemodynamic HbO/HbR responses using B-spline, we selected 10 knots pivoted at the percentiles of time sequence *t* = 0*s to* 30*s* with a step of 0.5*s*; therefore, 10 corresponding weights and basis functions, as illustrated in the second column of Fig. 3. Using this Bayesian spline model, not only the averaged time course of HbO/HbR, but also their corresponding standard deviation over the 101 sub-averaged, for each time point, were projected in this ‘spline space’. This means the averaged time course of HbO/HbR can be recovered by the linear combination of the mean of each weight (over 101 sub-averaged) and basis functions *B_n_*(*t*), whereas the standard deviation of HbO/HbR time course is reflected by the linear combination of the standard deviation of each weight (over 101 sub-averaged) and *B_n_*(*t*). Note that the use of spline basis functions in this study was mainly to reduce the dimensionality of the HbO/HbR time course from 60 sampling points to 10 weights while preserving the variability structure to be modeled. Therefore, selecting 10 spline knots was a trade-off: 1) choosing fewer knots that would result in distortions of the HbO/HbR time courses, involving too much temporal smoothness; 2) adding more knots would increase the dimensionality of the data after projection. Hence, our empirical choice allowed us to ensure an exact representation of the whole HbO/HbR time courses with a minimum dimensionality. Importantly, projecting to spline space also preserved the autocorrelation of HbO/HbR time courses per se, which could not be achieved when simply applying the same hierarchical model to each of the 60 data time points independently.

We then embedded this spline model of the hemodynamic response within the same hierarchical model proposed in the previous section (Model#1), this time replacing variables 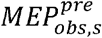 by the spline weights 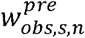 as follows, (see the third column of Fig.3).

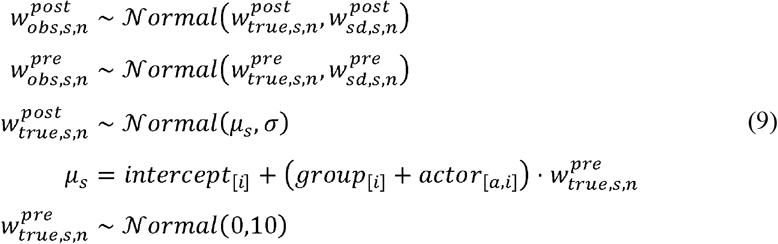

where 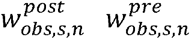, 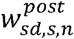 and 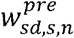 were all calculated from the corresponding posterior of spline weights estimated from equation (8). 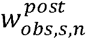 and 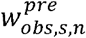 are referring to the mean of each spline weight for either pre- or post-PAS HbO/HbR for session *s*, estimated from the corresponding posterior of spline weights in equation (8). The scale of Gaussian distribution in the measurement error model was estimated as the standard deviation of the spline weights, 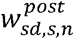 and 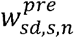 respectively.

Note that equation (8) resulted in the estimated posterior distribution of 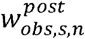 and 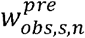 after projecting HbO or HbR time course to the spline space, the scales of Gaussian distributions used for the measurement error model in (9) were directly reflected by the standard deviation of the posterior distribution (denoted as *sd* in subscript). Finally, the intervention, subject index variables, and priors considered for this model, were similar to those previously introduced for Model#1, so the PAS effects on HbO/HbR whole time course were then encapsulated in the hierarchical model of spline weights.

#### 2.4.3 Hierarchical Bayesian Model #3: Relationship between PAS effects on task-related hemodynamic responses and PAS effects on cortical excitability

In this third model presented in Fig.4, we propose to investigate the interactions between 1) PAS effects on M1 excitability (PAS effects on MEP, represented by the slope parameter in Model#1), and 2) PAS effects on reconstructed hemodynamic finger tapping responses (PAS effects on HbO/HbR, represented by the slope parameter in Model#2 for a specific weight *w_n_*). We assumed the relationship between task-related hemodynamic responses and M1 excitability was not intervention specific, therefore, only the index variable of session *s* was used, whereas intervention and actor index *i* and *a* were ignored.

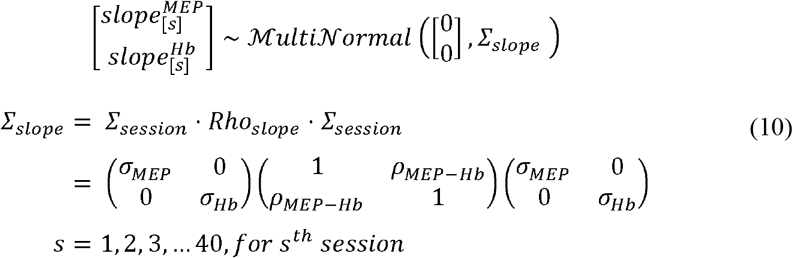

The interactions between task-related hemodynamic responses and cortical excitability were modeled using a multinormal distribution, in which the parameter *ρ_MEP–Hb_* in the *Rho_slope_* matrix denotes the correlation between the two slopes (representing PAS effects in both linear models). The same model was fitted separately when investigating either the relationship between MEP and HbO or between MEP and HbR. 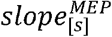 is the sessionspecific slope parameter in Model#1. Similarly, 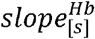 (either HbO or HbR) is the session-specific slope parameter in Model#2 for one of the corresponding spline weights *w*_1~10_. *σ_MEP_* and *σ_Hb_* are the standard deviations of 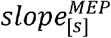 or 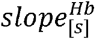, respectively. Note that this model was fitted for each spline weight separately – therefore we conducted *n* = 10 correlation investigations between MEPs and each spline weight for HbO and then for HbR. The posterior distribution of *ρ_MEP-Hb_* inferred for a specific spline weight can then be interpreted as the correlation between brain excitability and task-related hemodynamic responses at a specific time period. For instance, *w*_5_ reflects HbO/HbR fluctuations around the peak time point of the hemodynamic response. For the other parameters of the Multinormal distribution, we considered the same weakly informed priors as those proposed in Model#1 and Model#2.

**Fig.4.**
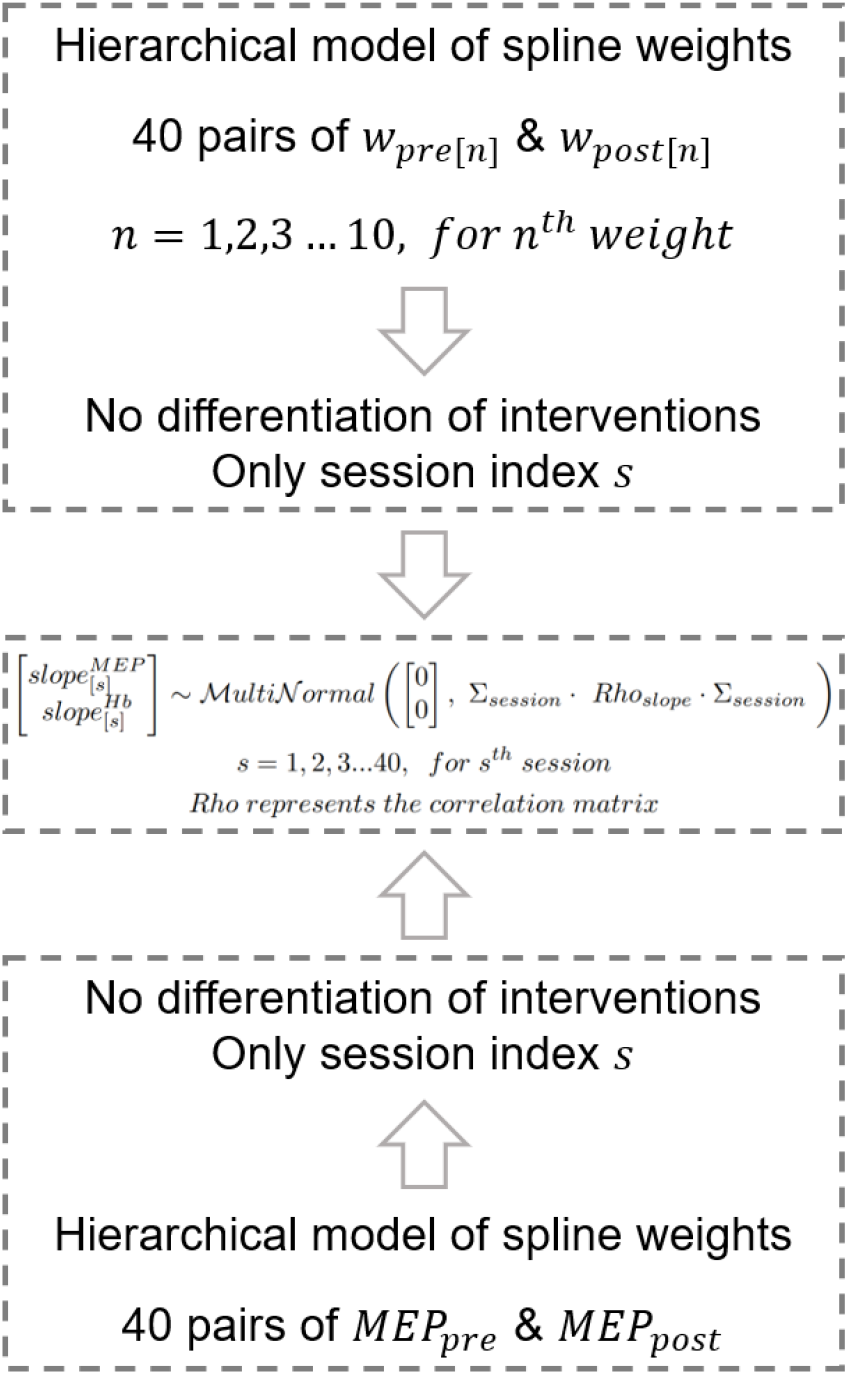
Modeling the relationship of task-related hemodynamic responses and cortical excitability. For MEP, which represented the M1 excitability, the previous Model #1 was modified to be session-specific only. For spline weights, which represented the features of task-related HbO/HbR time course, the previous Model #2 was also modified to be session-specific only. The association between the slope_MEP_ in MEP model and slope_Hb_ represented by any of the spline weight of the HbO/HbR time course model were described by a multinormal distribution.

### 2.5 Prior predictive simulation

To justify our choices of ‘weakly informed’ priors in Model #1, a prior predictive simulation was conducted and the corresponding results are presented in Fig. 5. The prior predictive simulation consists in a generative process simply checking what kind of data we would expect to generate from our hierarchical models, when applying all possible values of the parameters associated with the proposed prior distributions of the model, therefore assessing only the generative properties of our model. Then by comparing the distribution of data generated by our model, to the domain knowledge, one can assess whether the proposed priors could be overregulating or not objective (e.g., too strongly informed). In our study, PAS effects were modeled using linear regression. To perform prior predictive simulation, we draw 1000 lines following the prior distributions of the intercept and slope in the normalized pre-MEP vs. post-MEP amplitude plane. Then the distribution of generated regression lines was compared to three reference lines summarizing our knowledge of the problem. In further detail, these three reference lines were featuring a slope of 0.2, 1 and 3, respectively, and an intercept of 0. When the intercept is set to 0, the slope just refers to the ratio of post-over pre-PAS MEP amplitude, which was used in conventional analysis to represent the PAS effects [Cai et al., 2022b]. Whereas a slope of 1 would then correspond to no effect (post-/pre-PAS ratio of 1), the reference slopes of 0.2 and 3 represented the thresholds for outliers of extremely small or large MEP ratios [Kriváneková et al., 2013].

**Fig.5.**
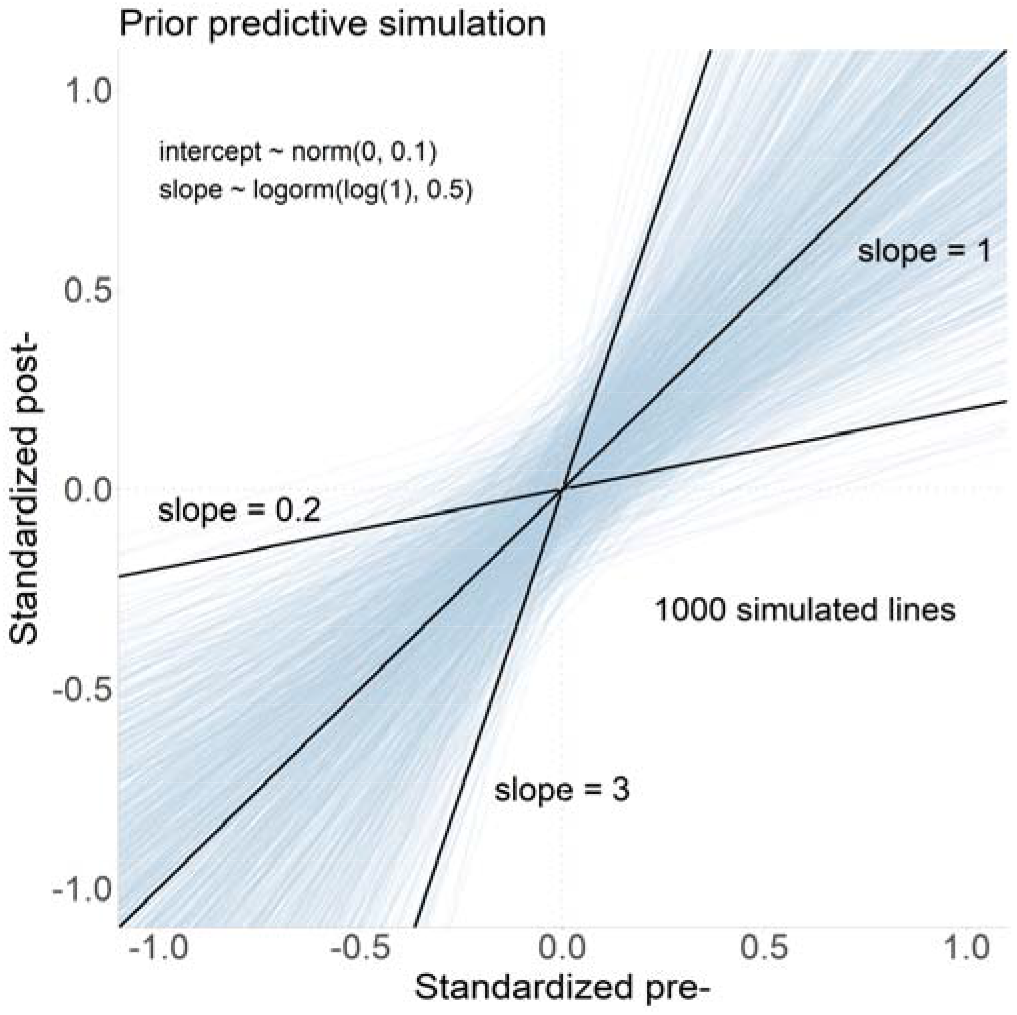
Prior predictive simulations for the hierarchical model of PAS effects on cortical excitability. Each blue line represents one prior predictive simulation obtained by drawing simultaneously the intercept and the slope parameters when considering only the priors proposed in Model#1. For comparison purposes, as a reference, we first represented a control line suggesting no PAS effect (intercept of 0, slope of 1), then two lines referring to MEP ratio outliers (intercept of 0 and slope of 0.2 and 3 respectively).

### 2.6 Hierarchical Bayesian model fitting

In this study, we used the R Version 4.0.3 [R Core Team, 2020] distribution of the Stan Probabilistic programming languages [Stan Development Team, 2020a] - RStan package Version 2.21.2 [Stan Development Team, 2020b] to implement and solve the proposed Bayesian models. Specifically, the joint posterior distribution was sampled using the implementation of dynamic HMC in Stan [Betancourt, 2017; Betancourt, 2019], as an improved version of HMC algorithm [Neal, 2010; Betancourt and Girolami, 2015]. In total, 4 MCMC chains were used to sample each model and they were initialized randomly to ensure a better exploration of the joint posterior distribution while allowing diagnosis of the convergence. Each chain consisted of 2000 samples, including a first half warm-up phase (1000 samples) for the adaptation of the HMC parameters. Therefore, when combining all 4 chains, we obtained 4000 samples of each parameter of the models mentioned above, drawn respectively to estimate the corresponding posterior distributions. Regarding computation time, using an Intel 10750H laptop CPU and parallel computation (one core per chain), dynamic HMC took 66s for sampling once Model#1, 59s for Model#2 and 53s for Model#3 (including compiling time and calculation of the diagnostics).

Diagnosing the HMC sampling process is a crucial step when evaluating the accuracy and biases of the estimated posterior distributions. This is also known as a unique and advanced feature of HMC when compared to other MCMC algorithms [Roberts and Rosenthal, 2004]. In this study, we considered the diagnostic approach recommended by Stan to evaluate pathological behaviors of HMC sampling [Betancourt, 2017; Gabry et al., 2019; Gelman et al., 2020b],

1. ***Divergent transitions for real samples were drawn after the warm-up phase.*** This diagnostic statistic is specific for the HMC sampler, mainly invigilating the miss-match between the step size of the MCMC chain and the target distribution geometries [Betancourt et al., 2017]. While sampling a ‘high curvature’ region of the target distribution, an inappropriate large step size may miss-sample it, therefore biasing the resulting posterior distribution. MCMC chains will approach infinite energy immediately – called divergent transitions – when approaching such regions [Neal, 2010; Betancourt, 2017]. These divergences are recorded and reported by Stan. Note that divergence is usually related to the parameterization of the model, especially when involving hierarchical structures. Parameters may usually be dependent on each other in these models, therefore, creating a ‘high curvature’ distribution landscape, also denoted as Neal’s Funnel [Neal, 2003], which is difficult to sample. In our study, in order to reduce the chances of such divergences, we considered the reparameterization of the model into non-centered forms when sampling with HMC.
2. ***The Energy-Bayesian Fraction of Missing Information (E-BFMI) is a specific diagnostic statistic for HMC sampler, evaluating the efficiency of the sampling process***[Betancourt, 2016]. Poorly chosen parameters of the HMC can decrease the efficiency of the sampling process or even result in incomplete exploration of the target distribution, especially when considering distributions with heavy tails. Such a behavior can be diagnosed by taking advantage of the physics feature of HMC, i.e., by comparing the marginal energy density (denoted as *π_E_*) and energy transition density (denoted as *π*_Δ*E*_) of the chain. When superimposing the histograms of *π_E_* and *π*_Δ*E*_, the higher the efficiency, the more overlap between the two distributions. The Energy Bayesian Fraction of Missing Information (E-BFMI) [Rubin, 2004] is used in Stan to quantify such comparison, by calculating the statistical expectation of the variance of *π*_Δ*E*_ over the variance of *π_E_*. Empirically, an E-BFMI value below 0.3 is considered as problematic [Betancourt, 2016; Betancourt, 2017].
3. ***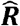 as a general and primary diagnostic statistic when evaluating convergence of MCMC chains*** [Gelman and Rubin, 1992; Brooks and Gelman, 1998]. 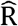 is estimated for each parameter of the model as the ratio of between-chains variance over the within-chain variance. In detail, the between-chains variance is calculated as the standard deviation among all chains, whereas the within-chain variance is calculated as the weighted sum of the root mean square of the standard deviation within every chain. The recommended criteria for convergence is 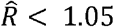 [Gabry et al., 2019; Vehtari et al., 2020].

Finally, we used tidyverse package [Wickham et al., 2019] in R [R Core Team, 2020] for general data wrangling and visualization. Tidybayes package [Kay, 2020] was used for visualizing the posterior distributions whereas bayesplot package [Gabry et al., 2019; Gabry and Mahr, 2020] was used for visualizing the diagnostics of HMC chains.

### 2.7 Statistical inferences of the models

We considered two types of statistical inferences in this study. To infer PAS effects on MEP and HbO/HbR time course, we first applied ***the posterior predictive simulation technique*** in which the distribution of MEP after each intervention type was estimated by feeding the fitted model (e.g., Model#1) with specific pre-intervention MEP values (the same approach was applied to HbO or HbR time courses in Model #2, respectively). Then, the distribution of percentage change of post-intervention MEP relative to pre-intervention MEP was used to infer the PAS effects on MEP (the same approach was applied to HbO or HbR time course, respectively). To assess the correlation between PAS effects on MEP and its effects on HbO/HbR, we directly considered the posterior distribution of the correlation parameter in Model#3. Please find further details as follows.

1. When investigating the effect of PAS on MEP, we answer the following question - what will be the distribution of MEP after a certain PAS intervention when providing a specific pre-PAS MEP amplitude as input? This approach is more direct and convenient compared to the process of checking the posterior distribution of each parameter of the model one by one. This technique is referred to as the posterior predictive simulation [Gabry et al., 2019; Gelman et al., 2020b]. For instance, ***to infer the PAS effects on the M1 cortical excitability,*** we used the averaged MEP (i.e., equal to 1.0mV, in the original data scale before normalizing) among all pre-PAS runs to represent the group-level pre-PAS M1 cortical excitability. This amplitude was then substituted into the fitted Model#1 along with all posterior distributions of parameters (e.g., intervention-specific intercepts and slopes) to estimate a group-level post-PAS MEP distribution. By comparing the distributions of the percentage change of this post-PAS MEP distribution relative to the pre-PAS MEP amplitude, the effects of each intervention can be inferred. We also performed this type of inference using a set of different pre-PAS MEPs values, such as 0.2mV, 0.6mV, 1.2mV, 2.2mV and 2.8mV according to the observed range of all individual pre-PAS MEP (i.e., ranging from 0.1mV to 3.0mV), therefore investigating how PAS effects could be related to the pre-PAS MEP amplitude. Note that these pre-PAS MEP amplitudes were also scaled by dividing the global maximum value of 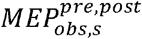 before being fed into the model. Finally, our resulting posterior predicted post-PAS MEP amplitudes were rescaled back to the original data scale. We considered a similar approach to statistically assess ***the PAS effects on task-related hemodynamic*** using the following steps: 1) select any preferred pre-PAS HbO/HbR time course (e.g., the averaged HbO/HbR of all pre-PAS runs demonstrated in the results); 2) calculate the 10 weights corresponding to this specific time course; 3) inferring the 10 post-weights along with their variance by posterior predictive simulations of the fitted hierarchical Model#2; 4) apply a linear combination of 10 post-weight and basis functions to obtain the distribution of post-PAS HbO/HbR whole time course. Note that we also calculated the PAS effects on HbO/HbR by contrasting post-PAS25 or post-PAS10 hemodynamic responses to the one obtained in post-sham condition, in order to provide inferences not biased by the control condition (i.e., sham intervention). To do so, we subtracted from the posterior predicted distributions of post-PAS25 HbO/HbR time courses (or post-PAS10) the posterior predicted post-sham HbO/HbR time course. Therefore, the final inference of PAS effects on post-PAS HbO/HbR whole time course can be considered as unbiased effects relative to the control condition (i.e., sham).
2. ***The correlation between M1 cortical excitability and task-related hemodynamic responses*** can be estimated by directly inferring the posterior distribution of the correlation parameter *ρ_MEP-Hb_* per se. Note that this correlation distribution was estimated for each spline weight separately, therefore, the resulting posteriors can be used indirectly to infer the excitability association for each specific time point of the HbO/HbR time course. For instance, the posterior distribution of the correlation between 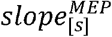 and 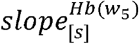 indicated the relationship between the peak period (e.g., *w*_5_ referring to a few seconds around the expected peak timing of the hemodynamic response) of task-related HbO/HbR and M1 cortical excitability. Moreover, we also conducted typical frequentist inferences of this relationship using the linear fit and Pearson’s correlation over all 40 sessions on the resulted mean of posterior distributions of 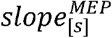 and 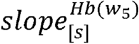 for both HbO and HbR.

Note that for summary statistics, we reported the median and the median absolute deviation (i.e., *mad_sd_*), which was suggested by Gelman et al., 2020a and estimated as follows: 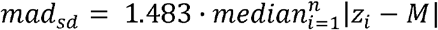, where *z_i_* is a certain value of a set of values *z*_*i*=1,2,3…*n*_ and *M* is the median of all *z_i_*. The *mad_sd_* is a more universal representation of the variance, which is comparable to the standard deviation, without considering the parametric/nonparametric distribution of *z_i_* and is more computationally stable.

### 2. 8 Data and code availability statements

The original raw data supporting the findings of this study are available upon reasonable request to the corresponding authors. fNIRS and TMS data were processed via Brainstorm software [Tadel et al., 2011] available at https://neuroimage.usc.edu/brainstorm/ and the fNIRS processing plugin - NIRSTORM (https://github.com/Nirstorm/nirstorm) in Brainstorm. R code for Bayesian models is available upon reasonable request to the corresponding authors.

## 3. Results

### 3.1 Prior predictive simulation

As illustrated in Fig.5, the resulting prior predictive simulation lines were distributed symmetrically around the control line suggesting no PAS effect (i.e., intercept = 0, slope =1). This means our priors exhibited no preference toward a slope < 1 or >1. Moreover, within the post-PAS MEP versus pre-PAS MEP plane, the area spanned by all simulated lines covered a larger area than the area enfolded by the reference lines, 0 intercepts, and the slope ranging from 0.2 to 3.0. These results confirm that the priors in our hierarchical model are not biased to the expected PAS effect and are more conservative than the conventional MEP ratio thresholding approach. Note that similar prior predictive simulation results could also be expected for PAS effects on HbO/HbR responses, since fNIRS responses were normalized similarly to MEP values and we also considered similar priors in the linear regression Models #1 and #2.

### 3.2 Diagnosis of HMC

All of the models considered in this study resulted in 0 divergences reported by Stan, indicating they were well parameterized, and HMC chains explored sufficiently well the target distribution [Betancourt, 2017; Gabry et al., 2019; Gelman et al., 2020b]. Fig.6 reports the evaluation of diagnostic statistics of 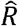 and E-BFMI. In each column of Fig.6, a specific model sampling process for a specific model is being diagnosed (see further details in Fig.6 caption). The first row illustrates the histogram of 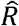 for all parameters in each corresponding model. No parameters resulted in 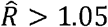 indicating the corresponding HMC chains converged properly. The second row demonstrated the superimposed histograms of *π_E_* (i.e., marginal energy density) and *π_ΔE_* (i.e., energy transition density), which overlapped well for all models. This evaluation was also quantified by reporting E-BFMI values for each model, which were all smaller than 0.3.

**Fig.6.**
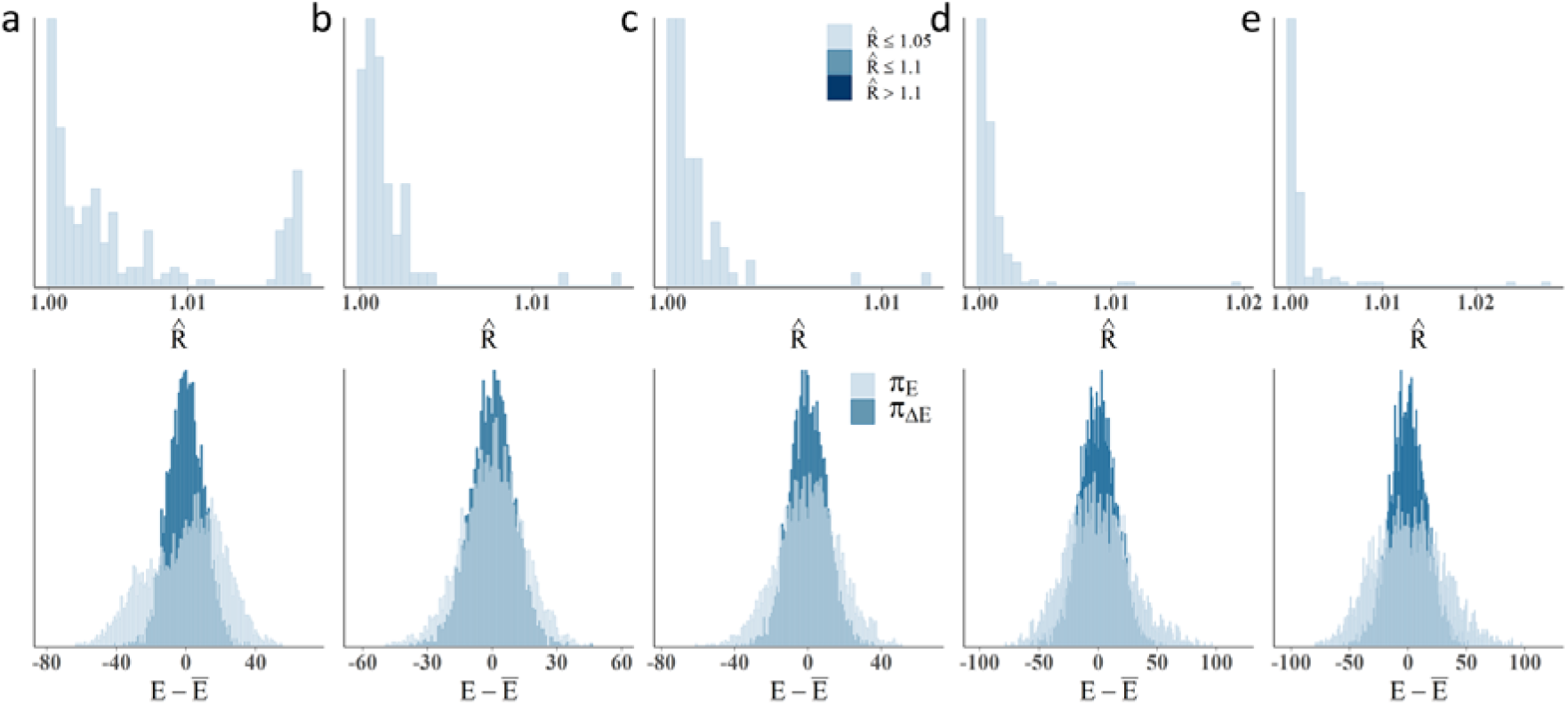
Diagnostic statistics of key features of the models considered in this study. Diagnostic statistics for a) Model#1 - PAS effects on MEP amplitude, b) Model#2 - PAS effects on *w*_5_ of task-evoked HbO, c) Model#2 - PAS effects on *w*_5_ of task-evoked HbR, d) Model#3 - correlation between PAS effects on MEP and PAS effects on *w*_5_ of task-evoked HbO and e) Model#3 - correlation between PAS effects on MEP and PAS effects on *w*_5_ of task-evoked HbR. The first row presents the histogram of 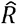 values for all parameters among all chains of each model. No 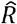 value was above 1.05, suggesting that all chains converged well. The second row presents the superimposed distributions of the marginal energy density π_E_ and the energy transition density π_ΔE_ for all HMC chains sampled for each model. E – Ē represents the centered energy. The corresponding quantification metric E-BFMI was smaller than 0.3, indicating a good overlapping between the two distributions. Similar results (not shown) were found for other weights.

### 3.3 PAS effects on cortical excitability

When considering Model#1, the estimated regression lines (using the averaged intercept and slope parameters calculated from their posterior distributions) linking pre- and post-PAS MEPs are reported in Fig.7a for each intervention. The regression line estimated for sham intervention (black line) was found, as expected, between the regression lines estimated for PAS25 (red line) and PAS10 (blue line), and it was almost identical to the reference line reporting no effect (intercept=0, slope=1). Observed pairs of post-PAS MEP and pre-PAS MEP mean amplitude over all trials are presented as solid points (observed data), whereas corresponding estimated ‘true’ amplitudes from posterior distributions are presented as empty points. The black lines connecting each pair of solid (observed mean) and empty (estimated ‘true’ mean) points illustrate the shrinkage, also known as the result of partial pooling obtained when considering hierarchical Bayesian modeling. This demonstrated the regularization property of the model, where the estimated ‘true’ MEPs corresponding to each intervention group shrank toward the corresponding regression line. Moreover, when considering the variance of the MEPs, the larger the MEP variability of a certain run, the more shrinkage there was.

**Fig.7.**
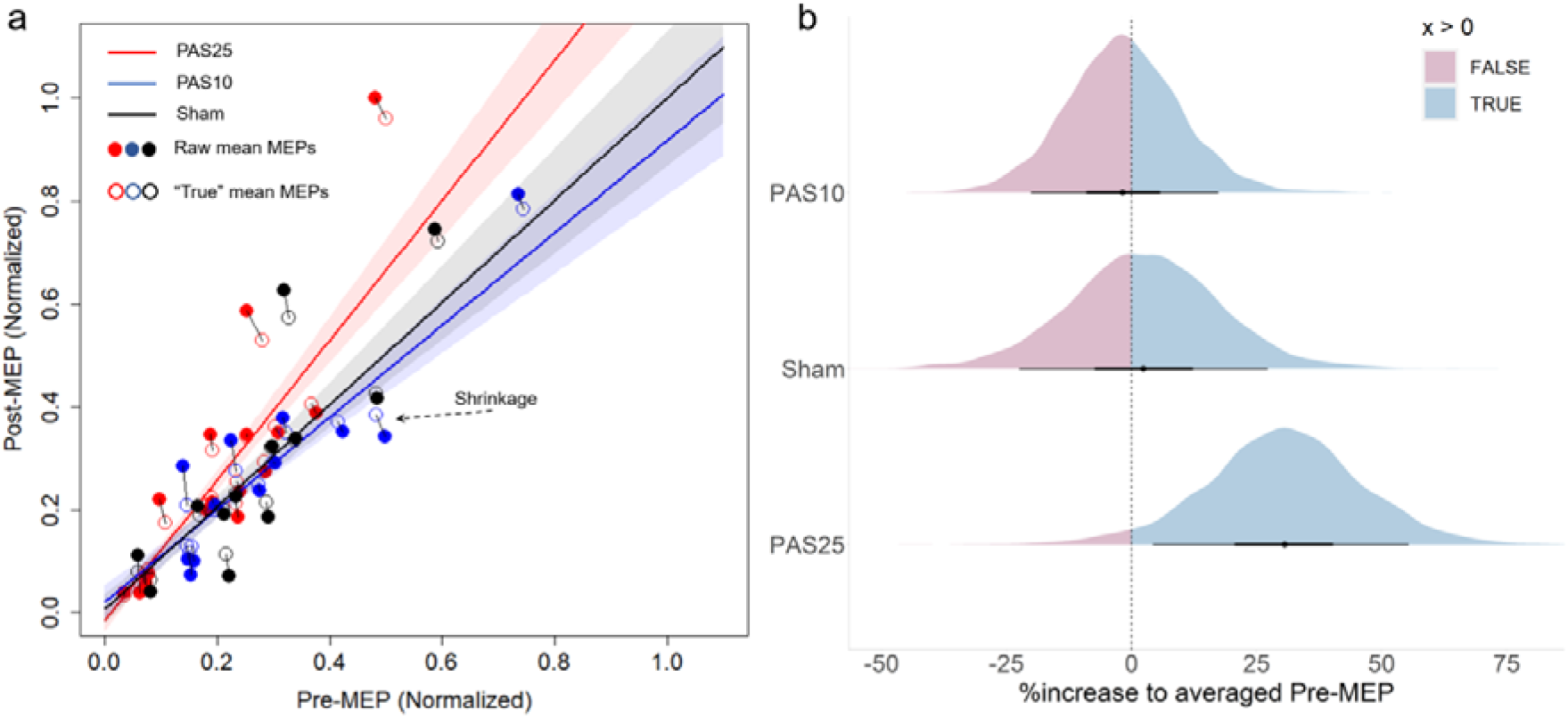
PAS effects on cortical excitability. a) the regression lines of each intervention estimated by the mean of intercept and slope from the corresponding posterior distribution, for PAS25 (red), PAS10 (blue) and sham (black). Pre- and post-PAS MEP amplitudes were normalized by dividing by the global maximum amplitude of all 80 MEP values. Shadow areas represent the 50% interval estimated from the posterior distribution of the regression parameters. Solid points correspond to pairs of averaged pre-/post-PAS MEP amplitudes over all trials of each specific run. Empty points represent the ‘true’ amplitude of the corresponding pre-/post-PAS MEP pair estimated using the proposed hierarchical Bayesian Model#1. The black bar connecting each solid point to the corresponding empty point illustrates the shrinkage process of Bayesian inference of the hierarchical model; b) Posterior predictive simulations of post-PAS MEP amplitudes obtained when considering a given pre-PAS MEP amplitude of 1mV as input, corresponding to the averaged pre-PAS MEP amplitude over all 40 sessions. The blue area represents the probability of obtaining a relative increase (in %) for the post-PAS MEP amplitude when compared to the pre-PAS MEP amplitude, whereas the pink area represents the probability of obtaining a relative decrease (in %). The black dot represents the median of each posterior distribution, and the surrounding horizontal lines show the corresponding 50% and 90% credibility intervals.

Results of posterior predictive simulation at the group-level are presented in Fig.7b. Considering a pre-PAS MEP amplitude of 1.0mV, the posterior distribution of relative changes of post-PAS MEP amplitudes (in %) after each intervention. PAS25 intervention resulted in a substantial relative increase in post-PAS MEP amplitude (*median±mad_sd_* = 30.6% ± 14.6%), consisting of a posterior probability of 0.97 for obtaining an increase in MEP amplitude. The posterior distribution of post-sham MEP amplitude exhibited a slight increase of 2.3% ± 14.5%. The effects of PAS10 were subtle, showing a slight shift towards the negative side consisting in a relative decrease of −1.80% ± 11.0%, and a probability of 0.57 of obtaining a decrease in MEP amplitude. Individual-level inferences are presented in Fig.S1, where both PAS25 and PAS10 effects are showing a relatively large between-subject variability.

Fig.8 presents the effects of simulating different pre-PAS MEP amplitudes as inputs, on the relative change of post-PAS MEP amplitude for each intervention, at the group level. For both PAS25 and PAS10, the higher the pre-PAS MEP amplitude was, the higher the relative change in MEP amplitude was. In further details, PAS25 resulted in an increase of post-PAS MEP amplitude of +26.2% ± 15.7% (*Prob* = 0.95), +31.4% ± 15.1% (*Prob* = 0.97), +33.5% ±17.7% (*Prob* = 0.96) and +33.9% ± 18.7% (*Prob* = 0.95) when considering an input pre-PAS MEP amplitude of 0.6mV, 1.2mV, 2.2mV and 2.8mV, respectively. Similarly, PAS10 resulted in an increase of post-PAS MEP amplitude of +4.2% + 18.9% (*Prob* = 0.59), when considering an input pre-PAS MEP amplitude of 0.6mV, followed respectively by decreases of –3.1% ± 10.5% (*Prob* = 0.62), –6.5% ± 12.6% (*Prob* = 0.70) and –7.51% ± 13.8% (*Prob* = 0.71) when considering an input pre-PAS MEP amplitude of 1.2mV, 2.2mV and 2.8mV. This important finding of our proposed Bayesian Model#1 suggests that even without increasing the TMS stimulation intensity during PAS, simply increasing the spTMS intensity considered to measure changes in excitability could have revealed the expected PAS effects more clearly, while reducing some variability in the data. On the other hand, when assessing this effect on sham, we obtained similar distributions of relative changes in post-PAS MEP amplitude, all symmetric around 0%, consisting in relative changes of 4.6%, 2.0%, 0.6% and 0.2%, when considering a pre-PAS MEP amplitude of 0.6mV, 1.2mV, 2.2mV and 2.8mV, respectively. Overall, when considering pre-PAS MEP amplitude of 0.2mV for each intervention, we found a large level of uncertainty in spTMS responses, suggesting that small MEP amplitude induced by spTMS should be avoided when assessing the level of brain excitability.

**Fig.8.**
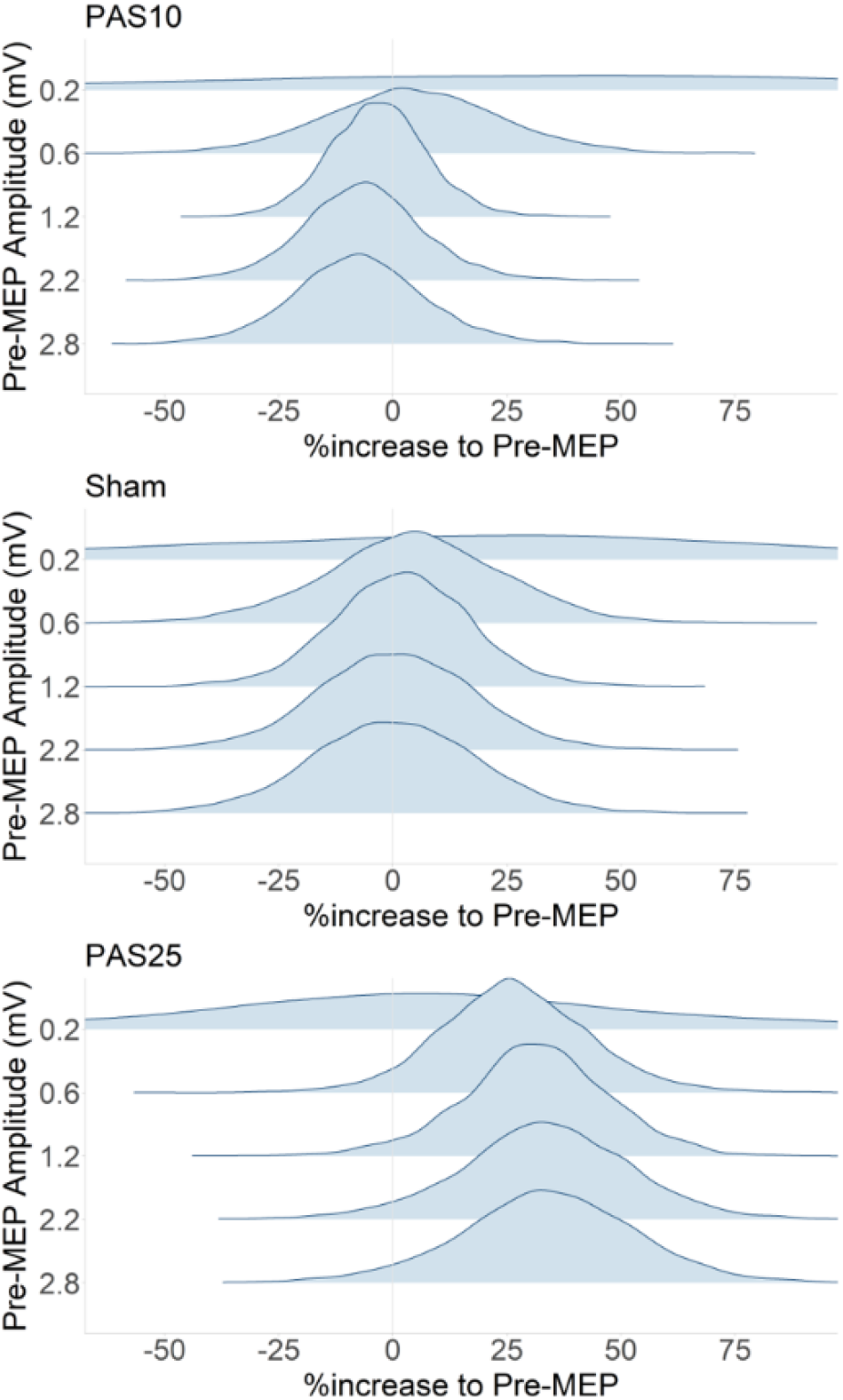
Effects of spTMS intensity on PAS assessment. We applied posterior predicting simulations when considering five levels of pre-PAS MEP amplitudes input, to evaluate the impact of five levels spTMS intensities. Posterior distributions of the corresponding relative changes in post-PAS MEP amplitude relative to pre-PAS MEP amplitudes are presented in each row. The expected effects of PAS25 (positive % increase) and PAS10 (negative % decrease) became clearer when increasing the spTMS intensity. On the other hand, when considering the sham intervention, we found no effect of relative changes in post-PAS MEP amplitude, for all intensity levels, whereas all posterior distribution remained symmetric at around 0%.

### 3.4 PAS effects on task-related hemodynamic responses

Fig.9 shows the PAS effects on the whole time course HbO/HbR. Results are reported here as a contrast between the intervention of interest (PAS25 or PAS10) and sham condition (i.e., subtraction of corresponding posterior distributions). When considering the group level averaged pre-PAS HbO/HbR responses (normalized to [-1, 1]) as input for posterior predictive simulations (dashed red and blue curves for HbO and HbR in Fig 9a), we found that PAS25 resulted in a relative increase of HbO amplitude (solid red curve) and negative decrease in HbR amplitude (solid blue curve), mainly around the expected peak of the hemodynamic response (from 8s to 16s, see Fig.9a). When comparing absolute peak amplitudes, the probability of increasing the hemodynamic response after PAS25 was 0.80 for HbO response and 0.82 for HbR response. After PAS10, our results at the group level suggested a subtle relative decrease of HbO and HbR absolute amplitudes around the peak of the hemodynamic response. The probability of obtaining a relative decrease in absolute peak amplitudes after PAS10 was 0.66 for HbO response and 0.48 for HbR response. Interestingly, PAS10 demonstrated a clear absolute amplitude decrease within a period ranging from the peak to the end of the response (11s to 25s) for both HbO and HbR. The aforementioned observations can be further illustrated by studying the posterior distribution of the response for specific time points along the window of the hemodynamic response. These results are reported in Fig9.b,c, assessing the hemodynamic response changes after PAS using posterior distributions for each selected time point. For instance, Fig.9 b and c illustrate the distributions of percentage changes in HbO (HbR respectively), at each time point (i.e., 4s, 8s, 10s, 12s, 14s and 18s) after PAS25 and PAS10, relative to the corresponding value at the same time but before the interventions.

**Fig.9.**
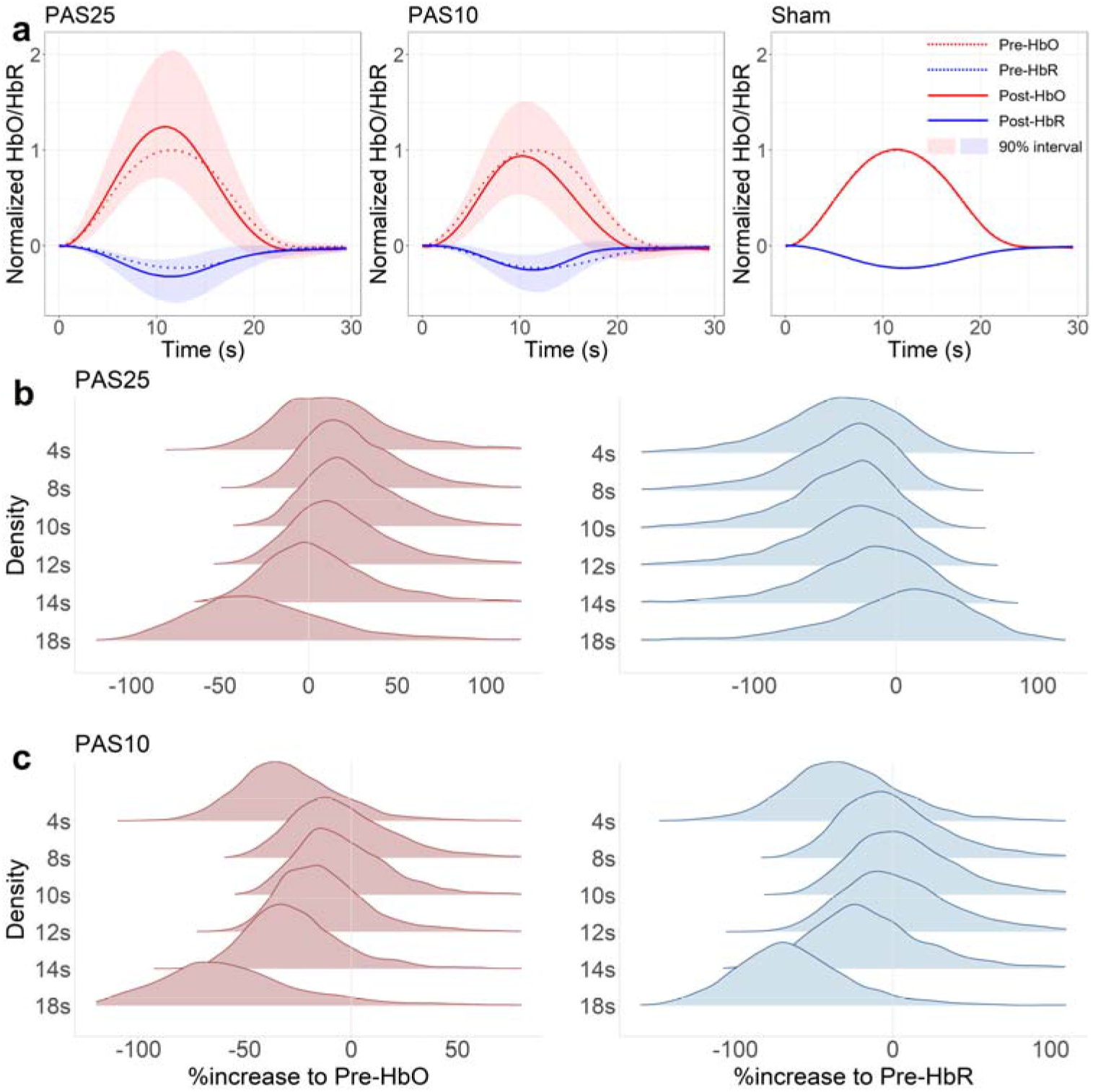
PAS effects on the whole time course of HbO/HbR. a) Posterior predictive simulations of post-PAS HbO/HbR time course (solid curves: HbO in red and HbR in blue) when considering pre-PAS HbO/HbR input defined as the group-level averaged pre-PAS HbO/HbR response (normalized to [-1,1]) over all 40 sessions (dashed curves: HbO in red and HbR in blue). The shadow area represents the 90% credibility interval of resulted post-PAS HbO/HbR responses. Note that sham effects were subtracted from PAS25 and PAS10 to obtain so-called ‘unbiased’ effects. The overlapping of curves (solid and dashed) in the sham panel are therefore shown as a sanity check of the contrast. b) Posterior distributions of the corresponding relative changes inferred at a set of different time points along the hemodynamic response (i.e., 4s, 8s, 10s, 12s, 14s and 18s). Post-PAS25 HbO amplitude relative to pre-PAS25 HbO amplitudes (red), post-PAS25 HbR amplitude relative to pre-PAS25 HbR amplitudes (blue). The x-axis is the % changes relative to the value (HbO or HbR) before PAS. c) Similar posterior distributions as in b) but for PAS10 results.

### 3.5 Relationship between PAS effects on task-related hemodynamic responses and PAS effects on cortical excitability

Inferences on the relationship between PAS effects on task-related hemodynamic responses and PAS effects on M1 excitability are presented in Fig.10a. We are reporting the posterior distribution of correlations between the slope of MEP amplitudes (post-PAS versus pre-PAS) and the slope of spline weight *w*_5_ (post-PAS versus pre-PAS) for either HbO or HbR task-related responses. Since our previous observations of the PAS effects were conducted for the whole HbO/HbR time course (Fig.9), we selected *w*_5_ as the spline weight of interest. Indeed, *w*_5_ spline corresponded to the basis function exhibiting a peak at 12.5 s, therefore consisting in the closest temporal pattern when compared to the expected hemodynamic response reported in Fig. 9. The probability of obtaining a positive correlation between PAS effects on MEP amplitude and PAS effects on HbO response was 0.77. The probability of obtaining a positive correlation between PAS effects on MEP amplitude and PAS effects on HbR response was 0.79. The corresponding 90% highest posterior density interval (HPDI) of this correlation was [-0.30, 0.88] for HbO; and [-0.22, 0.84] for HbR. Additionally, the linear fits between the posterior mean of 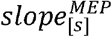 and the posterior mean of 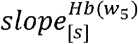 obtained for each session *s* among all 40 sessions is presented in Fig. 10b. The corresponding estimated Pearson’s correlation was 0.58 between MEP and HbO (p< .0001, CI_95%_ = [0.33, 0.75]) and 0.56 between MEP and HbR (p< .001, CI_95%_ = [0.30, 0.74]).

**Fig.10.**
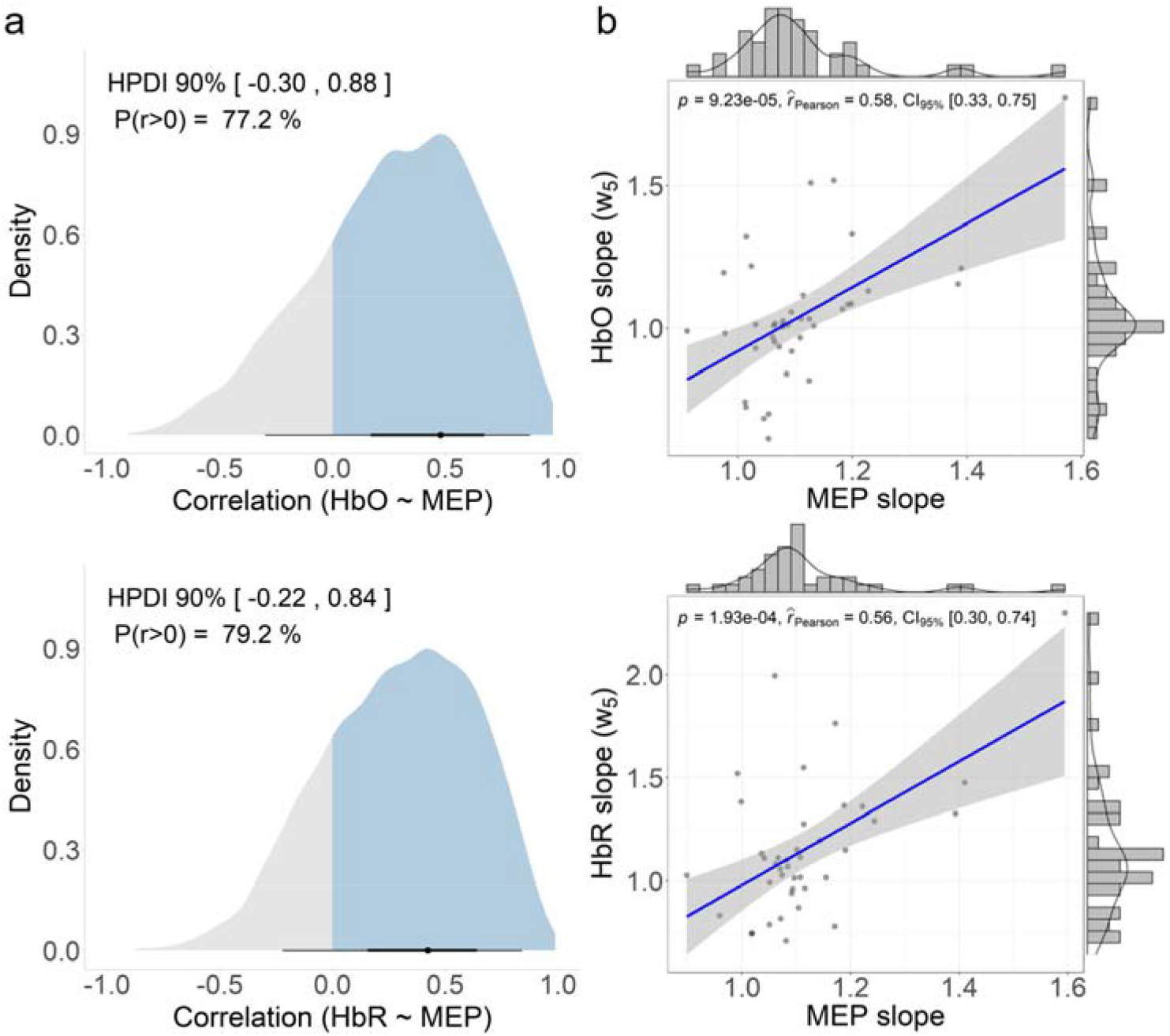
The relationship between PAS effects on task-related hemodynamic responses and PAS effects on cortical excitability. a) the posterior distribution of the correlation between 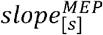 and 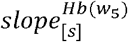 for HbO (top) and HbR (bottom). The blue area represents the probability of observing a positive correlation (ρ>0). The black dot represents the mode of each posterior distribution, and the surrounding lines show the corresponding 50% and 90% credibility intervals. b) Linear fit (blue line) between the posterior mean of 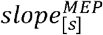 and the posterior mean of 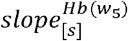 obtained for all 40 sessions (each represented by a grey dot). The grey area indicates 95% confidence interval of the regression. Estimated Pearson’s correlation 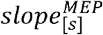 and 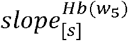 over the 40 sessions together with corresponding p-values and 95% confident intervals are shown on top of each panel. The marginal histograms and fitted density functions are shown on the side of each corresponding axis.

Fig. 11 illustrates the posterior distribution of the correlation between 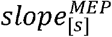 and 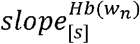 when considering each spline weight for *n* = 2,3,4,5,6,7 *and* 8 respectively. The closer the corresponding peak of the spline basis function associated with the weight *w_n_* was to the expected peak of the HbO/HbR response, the higher the correlation between and was. In further details, the median of these correlation values was respectively −0.05, 0.13, 0.28, 0.31, 0.13, −0.10 and 0.01 when considering for HbO; and a median value of 0.06, 0.17, 0.31, 0.32, 0.15, −0.06 and 0.03 when considering for HbR, therefore confirming this trend. Our results are suggesting that the expected positive correlation between PAS effects on task-related hemodynamic responses and PAS effects on M1 excitability appeared mostly around the peak of HbO/HbR time course (e.g.,), in agreement with PAS effects on hemodynamic responses, previously reported in Fig.9. On the other hand, for the earliest aspects of the hemodynamic response (modeled using) as well as for the end of the response (modeled using), we found a posterior correlation with a median close to zero, suggesting no relationship between and for the corresponding time periods of the hemodynamic response.

**Fig.11.**
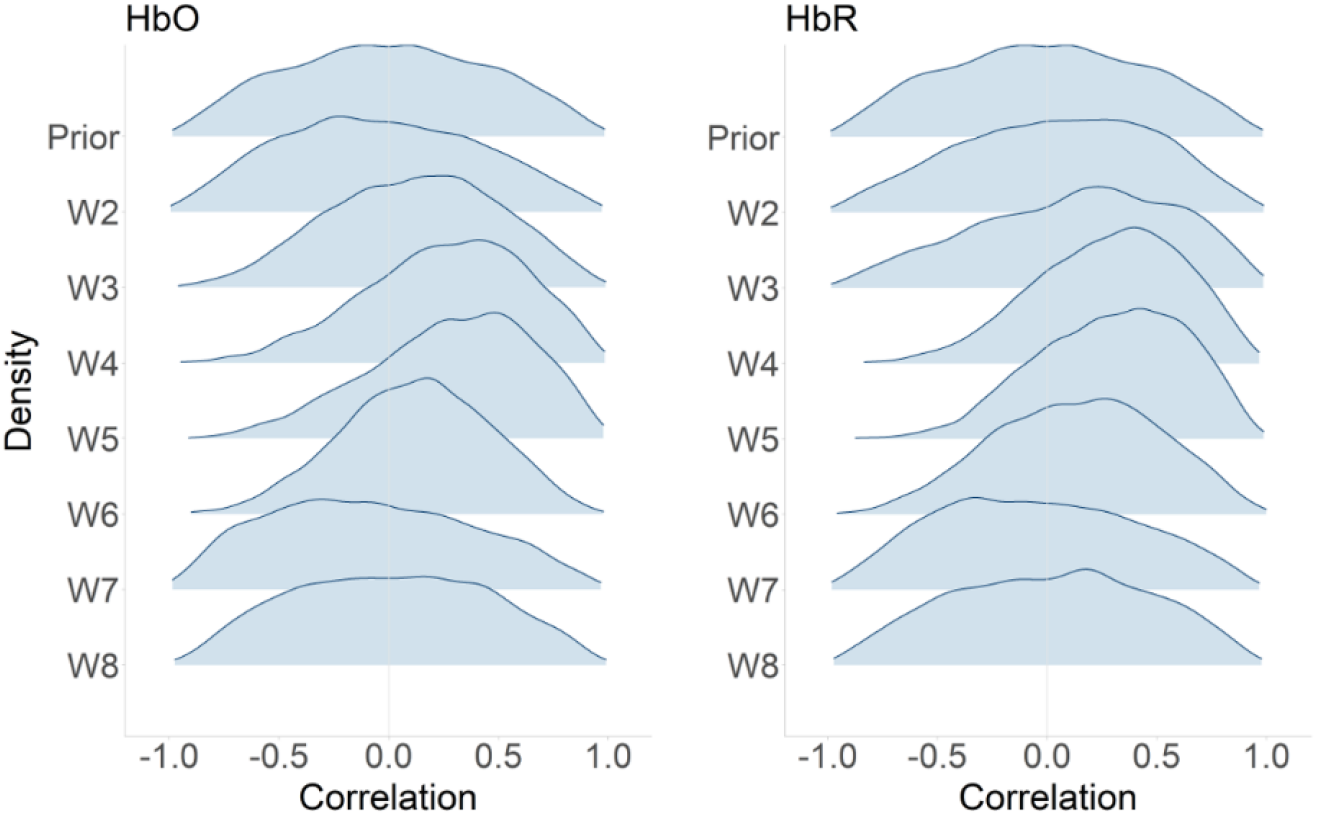
Posterior distributions of the correlations between and. The posterior distribution of the correlation between and, when considering each spline weights for for HbO (left) and for HbR (right). The prior distribution of the correlation value (i.e., LKJ(2) prior) is presented on the first row and is clearly exhibiting a symmetric distribution around the 0 correlation value. We found a trend suggesting that the closer the corresponding peak of the spline basis function associated with the weight was to the expected peak of the HbO/HbR response, the higher the correlation between and was. showed the highest correlation values for both HbO and HbR as it corresponded to the spline basis function exhibiting the peak at 12.5s, therefore the closest temporal pattern when compared to the expected hemodynamic response. In contrast, the earliest aspects of the hemodynamic response (modeled using w_2,3_) and the end of the response (modeled using w_7,8_), exhibited posterior distributions that are almost identical to the prior, therefore suggesting no effect.

## 4. Discussion

### 4.1 PAS effects on cortical excitability

Using hierarchical Bayesian modeling, we first investigated PAS effects on cortical excitability, which was measured using MEP amplitude induced by spTMS. Probability distributions of the relative changes (in %) of post-PAS MEP amplitudes, when compared to pre-PAS MEP amplitudes, were estimated using posterior predictive simulations. Our results showed a substantial increase in MEP amplitude after PAS25, a subtle decrease after PAS10, and a subtle increase after control (sham). These results are consistent with previous work performed with conventional analysis, calculating the ratio between the averaged MEP amplitude after PAS over the one before PAS [Stefan, 2000; Wolters et al., 2005; Tsang et al., 2015; Lee et al., 2017; Suppa et al., 2017]. Therefore, when the MEP ratio was larger than 1, it indicated an excitability increase and vice versa. In contrast, here we applied a full Bayesian workflow using an advanced sampling algorithm. There are several methodological advantages of our proposed procedure, as listed below: 1) linear regression allowed the differentiation of interventions and subjects, hence modeling the heterogeneity of intervention effects exhibited in different groups of data; 2) involving intercept in the linear regression reduced the influences of low MEP amplitudes runs when compared to the conventional ratio calculation of post-over pre-PAS MEPs; 3) the variability of MEP amplitudes were considered in the estimation of the PAS effects rather than only using the averaged amplitudes of each run and ignoring the variance; 4) parameters of the model were estimated by Bayesian inferences using dynamic HMC algorithm sampling posterior distributions using a hierarchical structure and weakly informed priors, therefore, allowing partial pooling to reduce the estimation uncertainty; 5) flexible and intuitive statistical inferences of the modeled PAS effects were obtained by conducting posterior predictive simulations from the model learned from the data. This means that by giving any pre-PAS MEP amplitude and intervention index, the distribution of the corresponding group-level post-PAS MEP amplitude could be estimated; Finally, 6) the estimated PAS effects were reliable and informative, as suggested by their posterior probability distributions, rather than considering only a statistical significance test providing a dichotomous output.

Moreover, our model also allowed assessing the effects of MEP amplitude itself on the effect size of the resulted excitability changes modulated by PAS. In Fig. 8, we reported a pattern suggesting that the higher the MEP amplitude was, the larger the effect size of both PAS25 and PAS10. This pattern was not biased when compared to the sham session, which showed no effects for different pre-PAS MEP amplitudes. Our results are in line with previous work on metaplasticity and state-dependence. We refer the reader to previous work on the topic for further details [Siebner et al., 2004; Silvanto and Pascual-Leone, 2008]. It is important to mention that when considering posterior predictive simulation, the intensity of the TMS pulse during the intervention session (i.e., during PAS25 or PAS10 or sham) did not change. This means that the underlying intervention effects did not change. Therefore, considering spTMS as the assessment procedure to measure brain excitability, our results are suggesting that a high enough spTMS intensity might help to measure more accurately and reliably PAS effects. Please also note that the pre-PAS MEP amplitudes (0.2mV, 0.6mV, 1.2mV, 2.2mV and 2.8mV) used in this posterior predictive simulation were selected based on the observed range of all individual pre-PAS MEP in our data (ranging from 0.1mV to 3.0mV). This observed range of MEP in our data from 16 subjects is also consistent with the spTMS evoked MEP distribution estimated by a recent meta-analysis study in which 687 healthy subjects’ data were considered among 35 studies [Corp et al., 2021]. More importantly, this meta-analysis study also investigated the relationship between the baseline MEP (referred to as the test stimulation) amplitude and the short interval intracortical inhibition (SICI) by pooling data from 15 studies consisting of 295 healthy subjects. They confirmed a significant negative relationship between the baseline MEP amplitude and SICI, suggesting that “SICI is best probed by high relative test stimulation intensities” [Corp et al., 2021]. This result is also concordant with our finding on PAS10, in which the higher the pre-PAS MEP (test stimulation) was, the more PAS10 was exhibiting clear inhibitory effects (decreased post-PAS MEP compared to the pre-PAS MEP). This consistency demonstrated the power of our Bayesian analysis to infer similar findings from relatively small sample-sized data when compared with meta-analysis results consisting of a much larger sample size.

### 4.2 PAS effects on the whole HbO/HbR time course of finger tapping response

To the best of our knowledge, this study demonstrated for the first time PAS effects on the whole time course of task-related HbO/HbR time courses. In contrast, only a few time segments within selected time windows were considered in previous studies, in which HbO or HbR amplitudes were just averaged at those selected time segments and compared before and after interventions [Chiang et al., 2007; Yamanaka et al., 2010]. In our previous study using conventional data analysis [Cai et al., 2022b], we also simply averaged the HbO/HbR amplitude within a 5s long time window centered around the peak of the hemodynamic response to represent the total amount of hemoglobin delivered to the region of interest. The Bayesian approach proposed in this study brings more insight into the investigation of PAS effects on hemodynamics, considering not only the peak amplitude before and after interventions, but whole HbO/HbR time courses. For instance, in Fig.9b and c, we can quantify the percentage change of HbO/HbR amplitude after either PAS25 or PAS10 interventions using their estimated probability distributions. Such quantification can be performed at any time point of interest (e.g., 4s, 8s, 10s, 12s, and 14s in Fig.9b and c). These results demonstrated the benefits of Bayesian approach when compared to conventional statistics - quantifying any effect of interest using its distribution. Visual inspections of results presented in Fig.9a suggest that PAS effects are indeed more pronounced around the peak of the expected hemodynamic response. This is expected if we can assume that the underlying hemodynamic response function (HRF) is not much affected by interventions. If so, the expected task-related hemodynamic response would result from a convolution with a higher or lower amplitude boxcar function representing the amount of excited or inhibited neuronal activity patterns [Sotero and Trujillo-Barreto, 2007]. Therefore, the effect of intervention should appear mostly around the peak, and the closer to the peak the higher the effect size. Consequently, averaging HbO or HbR amplitude within a certain time window would ‘dilute’ the estimation of the effect of interest, especially when considering the effect size was not large. For instance, we found an HbO increase of around 25% after PAS25 (see Fig.9a). Further averaging in time HbO response would reduce this percentage.

The fact that PAS intervention effects could be observed mainly around the peak of hemodynamic time courses may also explain the difficulty of investigating similar questions using fMRI. Indeed, a typical BOLD signal is sampled around 0.5Hz using standard fMRI sequences (as opposed to 10Hz in our fNIRS data). Such low temporal resolution may not be sufficient to sample well the effects around the peak and could possibly explain why no PAS effects were found on BOLD signal changes in the PAS and fMRI study reported by Kriváneková et al., 2013. Besides, depending on how well fMRI BOLD samples and the actual peak of the hemodynamic response were phased-locked, the mismatch between the time of BOLD signal sampling and the actual peak of the response may introduce some confounds, when comparing BOLD signal changes before and after PAS interventions.

Another benefit of modeling the whole HbO/HbR time course is the possibility to offer alternative interpretations of PAS effects. For instance, our results in Fig.9 showed a slight time shift for HbO after PAS25 and a larger one after PAS10. Indeed, after PAS10 intervention, the peak time of HbO response shifted from 12s in pre-intervention to 10s in post-intervention. The decrease of HbO amplitudes after PAS10 was also mainly exhibited from 11s to 25s of the response time courses. These observations may suggest a more complex underlying mechanism of the effect of neuronal plasticity on neurovascular coupling. Further analysis using the deconvolution technique [Machado et al., 2021] to estimate the underlying hemodynamic response function associated to these hemodynamic responses may help us to better investigate such a potential mechanism but this was beyond the scope of this study.

It is important to mention that we did not perform a specific analysis for every time sample of the hemodynamic response. We regularized and reduced the dimensionality of the problem by projecting HbO/HbR responses on B-splines, as temporal basis functions. Therefore, PAS effects on hemodynamic responses were modeled using only 10 spline weights instead of 60 data points, whereas the actual post-PAS HbO/HbR time courses could then be fully retrieved from the estimated weights and the spline basis functions. The choice of the number and locations of knots might have limited the ‘resolution’ of our proposed correlation analysis. More advanced Bayesian spline approaches have been proposed, such as the penalized spline (P-spline) [Eilers and Marx, 2010; Ventrucci and Rue, 2016], which introduces an extra prior to regularize the number of effective knots. Non-parametric time series modeling techniques were also proposed in this context, without assuming the location of the knots along the time course. For instance, Gaussian process regression [Neal, 1998] characterizes the time course, such as the hemodynamic response, as an unknown function. Samples of the time course are then drawn from a multinormal distribution providing a full covariance matrix of all time samples. Our analysis could benefit from these non-parametric approaches to avoid limitations associated with the choice of the knots, but this was beyond the scope of this study.

It is also worth mentioning that these results of PAS effects on the whole HbO/HbR time courses also benefit from accurate time courses estimated by our fNIRS reconstruction workflow [Cai et al., 2021; Cai et al., 2022a]. In this workflow, the fNIRS acquisition montage was personalized in order to maximize detection sensitivity to a targeted ROI. Meanwhile, the MEM framework adapted from our previous works in the context of electro-/magneto-encephalogram source imaging [Chowdhury et al., 2013; Chowdhury et al., 2016; Grova et al., 2016; Heers et al., 2016; Hedrich et al., 2017; Pellegrino et al., 2020; Abdallah et al., 2022] for conducting fNIRS reconstruction [Cai et al., 2021; Cai et al., 2022a] also ensured accurate estimation of HbO/HbR time courses from reconstructed spatiotemporal maps. For instance, delays between HbO and HbR peak times were around 1s (see Fig. 9), which is consistent with our previous findings [Cai et al., 2022b] and fNIRS literature [Jasdzewski et al., 2003; Steinbrink et al., 2006].

### 4.3 Relationship between PAS effects on task-related hemodynamic responses and PAS effects on cortical excitability

We finally investigated the relationship between PAS effects on task-related hemodynamic responses and PAS effects on cortical excitability along the whole HbO/HbR time course. Here, the benefits of the Bayesian approach consist of two aspects. First, the estimation of PAS effects on MEP and HbO/HbR was more accurate by taking advantage of the partial pooling feature of hierarchical Bayesian Model #1 and Model #2. Therefore, when we applied a conventional Pearson’s correlation analysis using the mean PAS effects on MEP and HbO/HbR (values on x- and y-axis in Fig.10b), summarized from their posteriors while ignoring the variance, we found statistically significant correlation values. This new analysis, applying Pearson’s correlation on estimated posterior mean, showed an improvement when compared to our previous inferences [Cai et al., 2022b], in which we reported nonsignificant correlations when considering a fully conventional analysis approach. Our new finding is therefore illustrating the importance of extensive handling of data variability to conduct more accurate statistical inference. Second, when considering the results shown in Fig. 10a, Bayesian analysis was more informative since we could estimate the whole posterior distribution of the correlation instead of providing just a single correlation value in Fig.10b. Therefore, the uncertainty of the correlation value was better estimated when compared to conventional analysis. This is often considered a unique advantage of Bayesian data analysis. Despite a rather small sample size and the large variability of PAS effects, the hierarchical Bayesian models demonstrated a high probability of positive correlations between MEP and hemodynamic slopes (around the peak of HbO and HbR responses). This finding is consistent with previous results reported in animal studies [Allen et al., 2007; Seewoo et al., 2019]. The reliability of our approach was further confirmed by the small correlation values between PAS effects on MEP and hemodynamic responses, when considering other time windows, such as the start and the end of the hemodynamic response (Fig.11).

### 4.4 HMC sampling and diagnostic

Taking advantage of dynamic HMC to sample the hierarchical Bayesian models in this study, we were able to carefully diagnose the pathological behavior of MCMC sampling chains [Betancourt, 2017; Betancourt, 2019]. This diagnostic procedure is an essential step when applying Bayesian data analysis [Gelman et al., 2020b]. To allow accurate and reliable inferences, MCMC chains must explore well the typical set of the posterior distributions, in which most of the probability density is contained. For instance, the convergence of MCMC chains needs to be confirmed and quantified to ensure full explorations of posterior distributions. When inappropriate parameters of the chain are chosen (e.g., the step size), abnormalities such as divergences should be detected to avoid eventual sampling biases. In our study, we reported several diagnostic statistics for all key components of three models using both visualization and quantified metrics, following the recommendations of the Stan team [Gelman et al., 2013; Gabry et al., 2019; Stan Development Team, 2020a]. The proposed diagnostic statistics considered here also constitute a unique feature of HMC sampling, when compared to conventional MCMC algorithms such as Gibbs sampling [Geman and Geman, 1984; Gelfand and Smith, 1990]. HMC is also considered to be more accurate by taking advantage of sampling all parameters at the same time, compared to Gibbs sampling in which parameters are sampled alternatively one after the other which may bias the resulting posterior distribution due to the inherent correlations between parameters. Overall, the diagnostic analysis of the sampling process in this study is suggesting that our inferences are built upon well-sampled posterior distributions. Similar HMC sampling and diagnostic approaches were also reported in several recent studies, such as a Bayesian virtual epileptic patient to model the spread of epileptic activity [Hashemi et al., 2020]; a Bayesian latent spatial model for mapping biomarkers of the progression of Alzheimer’s disease [Dai et al., 2021]; the Bayesian multilevel modeling to improve statistical inferences in fMRI analysis [Chen et al., 2019b; Chen et al., 2019a; Chen et al., 2021] and a hierarchical Bayesian model to investigate mechanisms of reinforcement learning and decision-making [Ahn et al., 2017].

### 4.5 Limitations and perspectives

We only involved one model for each investigation in this study. It is indeed recommended to construct multiple models based on different hypotheses of the same question and then quantitatively compare these models using techniques such as cross-validation to choose the most reliable one, providing a trade-off between overfitting and underfitting [Gelman et al., 2020b]. For instance, we proposed a linear relationship between cortical excitability and hemodynamic responses evoked by a finger-tapping task. However, such association might reach a plateau when excitability changes are either too low or too high, suggesting some non-linear effects. Moreover, the neurovascular coupling process includes different aspects like excitatory and inhibitory neurons, glial cells, the vasculature components like pericytes [Iadecola, 2017]. The interaction between inhibitory and excitatory neurons, the glial cell mediated signaling pathways, and their role in neurovascular coupling has been highly simplified in this linear model. A more detailed metabolism model involving blood flow dynamics [Buxton, 2021] may improve our inferences by comparing it with the model proposed in this study. Considering such advanced model comparisons, applied within a Bayesian framework, could be of great interest but was out of the scope of this study. Moreover, we conducted TMS following the recommendations of the International Federation of Clinical Neurophysiology (Rossi et al., 2009), which means our data set should not explore extreme conditions between excitability and hemodynamic responses, which are more likely to exhibit eventual nonlinear relationships.

Another limitation of our study was that M1 excitability was not assessed at the same time as the finger-tapping task, but sequentially, hence we could not apply a fusion model to pool the relationship between cortical excitability and hemodynamic responses at the single-trial level. We considered the mean and variance of MEP amplitudes and HbO/HbR time courses within the whole session as the input for the correlation analysis. This might reduce the resulted correlation values considering additional fluctuations of the baseline excitability and hemodynamic responses. However, since it has been shown that PAS modulated excitability changes could last for more than 30 minutes [Stefan, 2000; Lee et al., 2017], we are confident that our investigation of cortical excitability using MEP after spTMS and hemodynamic responses elicited by finger tapping was indeed still within this PAS effective duration window.

As perspectives for this study, it would be of great interest to investigate the relationship between spTMS evoked HbO/HbR and the corresponding MEP amplitude when occurring exactly at the same time, therefore, preventing confounds introduced by fluctuations of excitability and hemodynamic responses along the time. Such an investigation may help us in understanding the integrity of neurovascular coupling during transient cortical excitability changes induced by spTMS. Furthermore, the effect of stable cortical excitability changes (induced by PAS) on this integrity can be explored by comparing the spTMS evoked hemodynamic responses before and after PAS interventions. Additionally, since fNIRS data were recorded during the whole experiments (i.e., also during spTMS and PAS intervention), our data would allow assessing dynamically the evolution of MEP and hemodynamic responses during PAS. Such analysis would require modeling fNIRS response using advanced deconvolution techniques to handle the overlapping of TMS pulses induced hemodynamic responses [Machado et al., 2021], and will be considered in our future investigations.

## 5. Conclusion

In this study, we proposed hierarchical Bayesian modeling to investigate the relationship between motor task-related hemodynamic responses and M1 excitability. When compared with a sham control condition, a substantial M1 excitability increase was found after PAS25, and a subtle reduction of M1 excitability was found after PAS10. PAS effects on motor task-related hemodynamic responses were observed mainly around the peak of HbO/HbR time courses. We showed a high probability of positive correlations between PAS effects on MEP amplitudes and hemodynamic responses. Such correlations were also mainly exhibited around the peak of HbO/HbR time courses. Diagnostics of sampling MCMC chains showed no pathological behavior, ensuring the reliability of our results. Finally, this study also demonstrated the power of the Bayesian data analysis dealing with relatively high variability and small sample size data while providing informative inferences.

## Acknowledgments

This work was supported by the Natural Sciences and Engineering Research Council of Canada Discovery Grant Program (CG and JML), an operating grant from the Canadian Institutes for Health Research (CIHR MOP 133619 (CG)), a FRQNT research team grant, and a FRQS-Quebec Bio-Imaging Network (QBIN) Pilot Project grant. fNIRS equipment was acquired using grants from NSERC Research Tools and Instrumentation Program and the Canadian Foundation for Innovation (CG). ZC is funded by the Fonds de recherche du Québec – Sante (FRQS) Doctoral Training Scholarship and the PERFORM Graduate Scholarship in Preventive Health Research. GP is funded by Strauss Canada Foundation and a McGill/MNI - Fred Andermann EEG and Epilepsy Fellowship.

## Conflict of interest

The authors declare no potential conflict of interest.

## Appendices

## Appendix 1 fNIRS data processing

In this section, we are describing in further detail our fNIRS data analysis workflow. Raw fNIRS data were first preprocessed following standard recommendations [Yücel et al., 2021]. a) bad channel rejections of channels exhibiting either a negative raw amplitude during the whole time course and a coefficient of variation (CV) larger than 8% [Schmitz et al., 2005; Schneider et al., 2011; Eggebrecht et al., 2012; Piper et al., 2014]: b) linear regression of superficial physiological fluctuations using the average of all proximity channels [Zeff et al., 2007]; c) band-pass filtering (i.e., 0.01Hz to 0.1Hz) using a 3rd order Butterworth filter (zerophase); d) conversion in optical density changes (i.e., ΔOD) using logarithm conversion; e) ΔOD epochs extraction within a time window ranging from −10s to 30s around task onsets. Instead of the conventional process of averaging extracted ΔOD epochs, we then conducted a resampling process to estimate not one but a set of ‘possible’ averaged ΔODs [Cai et al., 2022b]. Our rationale was to propose an evaluation preserving the intrinsic variance of averaged ΔOD related to the underlying physiological fluctuations and eventual measurement errors such as motion artifacts. To do so, we first averaged 16 out of 20 preprocessed ΔOD epochs for all possible unique combinations (i.e., 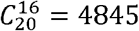 possibilities). Then, the averaged signal to noise ratio (SNR) of the resulting averaged ΔODs, for each wavelength, was estimated as the peak amplitude over the averaged standard deviation of baseline (within −10s to 0s) among all channels. Lastly, we selected 101 of these resampled averaged ΔODs, distributed around the median SNR (50 averaged below and 50 averaged above the median SNR), to obtain a distribution of ‘possible’ responses evoked by one finger-tapping run. The selection of 16 blocks out of 20 trials and 101 resampled averaged ΔODs maintained a good coverage of the data distribution. This number was empirically defined according to the observation that usually there were less than four blocks contaminated with artifacts in one finger-tapping run. Selecting sub-averaged trials around the median SNR ensured a good representation of fNIRS responses while discarding artifacts in the meantime. Indeed, in artifacts contaminated data, large motion artifacts would result in high SNR of corresponding sub-averaged trials.

We then applied 3D fNIRS reconstruction workflow using personalized optimal montage and maximum entropy on the mean (MEM), as further described and validated in our earlier work [Cai et al., 2021; Cai et al., 2022a; Cai et al., 2022b], to the 101 sub-averaged ΔODs. Therefore, ‘all possible’ HbO/HbR responses for each finger-tapping run were reconstructed as spatiotemporal maps along the cortical surface. To do so, the subject-specific fNIRS forward model was first estimated according to the following steps: a) 5 tissues head segmentation (e.g., scalp, skull, Cerebrospinal fluid, grey matter and white matter) calculated using FreeSurfer6.0 [Fischl et al., 2002] (https://surfer.nmr.mgh.harvard.edu) and SPM12 [Penny et al., 2011] (https://www.fil.ion.ucl.ac.uk/spm/software/spm12/); b) light fluences at each optode location, and for each wavelength (i.e., 685nm and 830nm), were calculated by simulating 10^8^ photons, using MCXLAB toolbox - a Monte Carlo photon simulator for modeling light transport in 3D turbid media, developed by Fang and Boas, 2009 and Yu et al., 2018; c) sensitivity of each voxel was computed using the adjoint formulation and was normalized by Rytov approximation [Arridge, 1999]; d) surface space sensitivity was finally obtained by projecting volumetric sensitivity map to subject’s cortical surface (i.e., midsurface, a middle layer of the gray matter, between pia mater and gray-white matter interface, 25,000 vertices) using the Voronoi based method proposed by Grova et al., 2006. Finally, each 101 averaged ΔOD epoch was down-sampled to 2Hz and MEM method proposed previously by our group for fNIRS reconstruction [Cai et al., 2021; Cai et al., 2022a] was applied to estimate the HbO/HbR spatiotemporal maps (0s to 30s) along the subject-specific cortical surface.

## Supplementary material

**Fig.S1.**
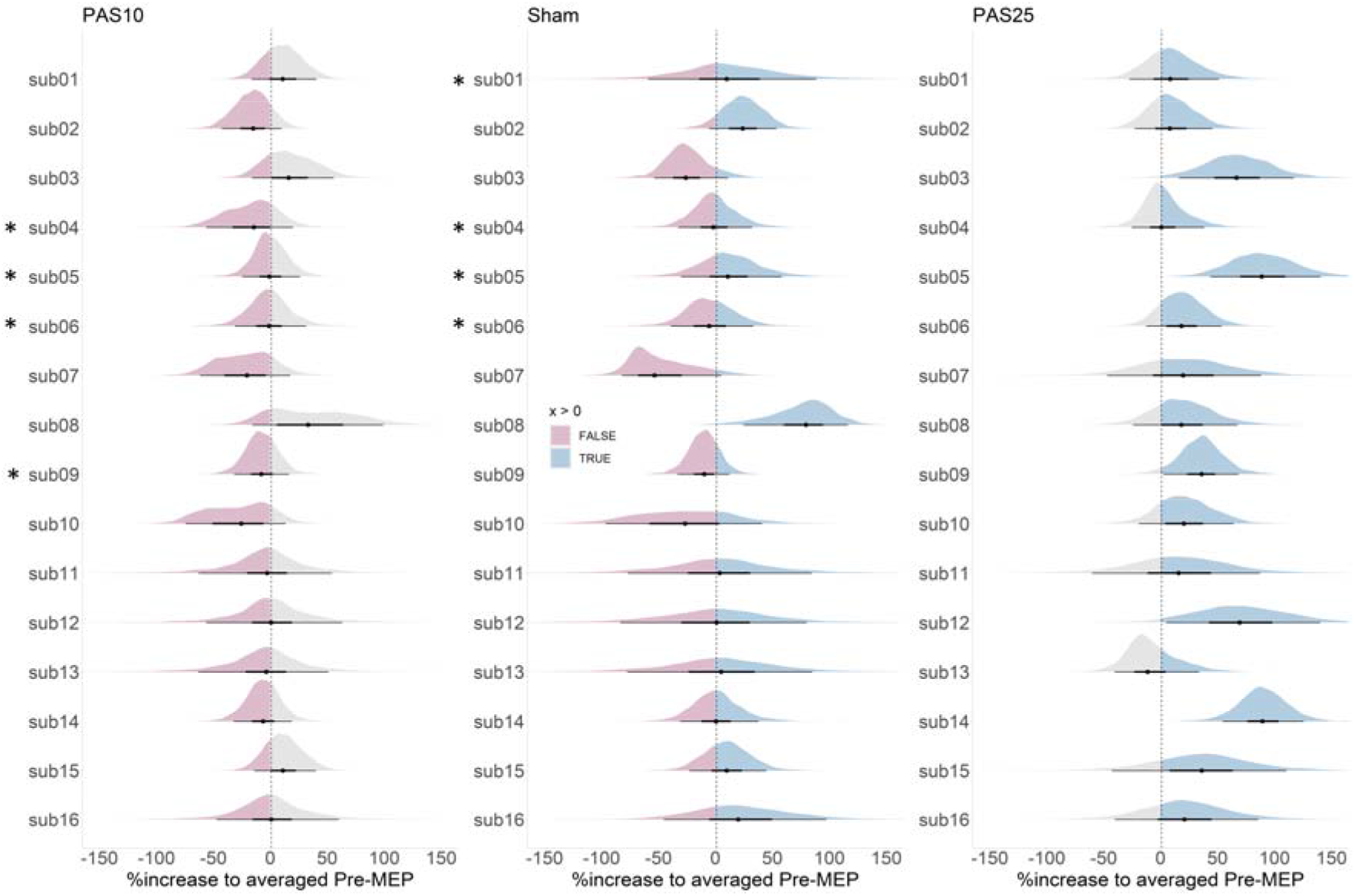
PAS effects on M1 excitability at the individual level. Each column presents the individual level posterior distribution of the intervention effect from PAS10, sham and PAS25, respectively. Posterior predictive simulations of post-PAS MEP amplitudes were conducted by assuming the same pre-PAS MEP amplitude for all subjects, i.e., the averaged pre-PAS MEP amplitude obtained for all subjects over all 40 sessions. The blue area represents the probability of obtaining a relative increase (in%) for the post-PAS MEP amplitude when compared to the pre-PAS MEP amplitude, whereas the pink area represents the probability of obtaining a relative decrease (in %). The black dot represents the median of each posterior distribution, and the surrounding bars show the corresponding 50% and 90% credibility intervals. Overall, large between-subject variability can be observed for both interventions. Note that missing sessions were also included using posterior predictive simulations within the model, based on prior distributions and partial pooled information from other sessions. * indicates the missing runs that were imputed by the model.

## Notes

### Competing Interest Statement

The authors have declared no competing interest.

### Summary of Updates

Revision

